# Genomic Abelian Finite Groups

**DOI:** 10.1101/2021.06.01.446543

**Authors:** Robersy Sanchez, Jesús Barreto

**Affiliations:** Department of Biology. Pennsylvania State University, University Park, PA 16802.; Universidad Central “Marta Abreu” de Las Villas. Santa Clara. Cuba.

**Keywords:** Genomics, Genetic code, Abelian groups, genome algebra, automorphism, mutational event

## Abstract

Experimental studies reveal that genome architecture splits into DNA sequence domains suggesting a well-structured genomic architecture, where, for each species, genome populations are integrated by individual mutational variants. Herein, we show that, consistent with the fundamental theorem of Abelian finite groups, the architecture of population genomes from the same or closed related species can be quantitatively represented in terms of the direct sum of homocyclic Abelian groups of prime-power order defined on the genetic code and on the set of DNA bases, where populations can be stratified into subpopulations with the same canonical decomposition into *p*-groups. Through concrete examples we show that the architectures of current annotated genomic regions including (but not limited to) transcription factors binding-motif, promoter regulatory boxes, exon and intron arrangement associated to gene splicing are subjects for feasible modeling as decomposable Abelian *p*-groups. Moreover, we show that the epigenomic variations induced by diseases or environmental changes also can be represented as an Abelian group decomposable into homocyclic Abelian *p*-groups. The nexus between the direct sum of homocycle Abelian *p*-groups and the endomorphism ring paved the ways to unveil unsuspected stochastic-deterministic logical propositions ruling the ensemble of genomic regions. Our study aims to set the basis for concrete applications of the theory in computational biology and bioinformatics. Consistently with this goal, a computational tool designed for the analysis of fixed mutational events in gene/genome populations represented as endomorphisms and automorphisms is provided. Results suggest that complex local architectures and evolutionary features no evident through the direct experimentation can be unveiled through the analysis of the endomorphism ring and the subsequent application of machine learning approaches for the identification of stochastic-deterministic logical rules (reflecting the evolutionary pressure on the region) constraining the set of possible mutational events (represented as homomorphisms) and the evolutionary paths.

## 1 Introduction

The analysis of the *genome architecture* is one of biggest challenges for the current and future genomics. Herein, with the term *genome architecture* we are adopting the definition given by Koonin [1]: *Genome architecture can be defined as the totality of non-random arrangements of functional elements (genes, regulatory regions, etc.) in the genome*.

Current bioinformatic tools make possible faster genome annotation process (identification of locations for genes, regulatory regions, intron-exon boundaries, repeats, etc.) than some years ago [2]. Current experimental genomic studies suggest that genome architectures must obey specific mathematical biophysics rules [3–6]. Experimental results points to an injective relationship: *DNA sequence* → *3D chromatin architecture* [3,4,6], and failures of DNA repair mechanisms in preserving the integrity of the DNA sequences lead to dysfunctional genomic rearrangements which frequently are reported in several diseases [5]. Hence, ***some hierarchical logic is inherent to the genetic information system that makes it feasible for mathematical studies*.** In particular, there exist mathematical biology reasons to analyze the genetic information system as a communication system [7–10].

We propose the study of genome architecture in the context of population genomics, where all the variability constrained by the evolutionary pressure is expressed. Although the random nature of the mutational process, only a small fraction of mutations is fixed in genomic populations. In particular, fixation events, ultimately guided by random genetic drift and positive selection are constrained by the genetic code, which permits a probabilistic estimation of the evolutionary mutational cost by simulating the evolutionary process as an optimization process with genetic algorithms [11].

### 1.1 The genetic code

Under the assumption that current forms of life evolved from simple primordial cells with very simple genomic structure and robust coding apparatus, the genetic code is a fundamental link to the primeval form of live, which played an essential role on the primordial architecture. The genetic code is the cornerstone of live on earth, the fundamental communication code from the genetic information system [8,9]. The code-words from the genetic code are given in the alphabet of four DNA bases 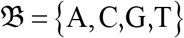and integrates a set of 64 DNA base-triplets {*XYZ*} also named *codons*, where 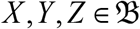. Each codon encodes the information for one aminoacids and each aminoacid is encoded by one or more codons. Hence, at biomolecular level, the genetic code constitute a set of biochemical rules (mathematically expressed as an injective mapping: *codon* → *aminoacid*) used by living cells to translate information encoded within genetic material into proteins, which sets the basis for our understanding of the mathematical logic inherent to the genetic information system [9,12].

The subjacent idea to impose a group structure on the set of codons resides on that the genetic code is the code of a communication system, the genetic information system [8,13,14]. As suggested by Andrews and Boss [15]: “*In codes used for electrical transmission of engineering signals, **group structure is imposed to increase efficiency and reduce error**. Similarly, the group characteristics of codon redundancy could serve to transmit additional information superimposed on the messages directing amino acid order in protein synthesis*”. As in the current human communication systems [16], to impose a group structure (on biophysical basis) on the set of codons facilitate a better understanding and evaluation of the error performance and efficiency of the genetic message carried in the chromosomes across generations [15].

### 1.2 The genetic code algebraic structures

The basis of the current study are algebraic structures (specifically groups structures) defined on the set of bases and on the codon sets. We assume that readers are familiar with algebraic structures like group, ring, and the classical mapping defined on them, homomorphisms, automorphism, and translations. For readers not familiar with this subject, a brief basic introduction to these definitions is given in the Appendix and the subject is well covered in textbooks elsewhere.

#### The meaning of group operations

Group operations are defined on the sets of DNA bases and codons, are associated to physicochemical or/and biophysical relationships between DNA bases and between codons and aminoacids. In other words, a proper definition of a group operation on the set of bases or on the set of codons will encode the physicochemical or/and biophysical relationships between the set’s elements. Thus, by group operations defined on the set of bases or on the set of codons, we understand an *encoding* applied to represent specified physicochemical or/and biophysical relationships as group operations between the elements of the set. Then, we shall say that such an encoding permits the *representation* of DNA bases, codons, genes, and genomic sequences as elements from an algebraic structure.

The encoding of physicochemical or/and biophysical relationships between DNA bases and between codons and aminoacids implicitly assumes that the genetic code is the code of a communication system; in the current case, the genetic information system [8,9]. It has been shown that the genetic code is robust to translation error and it is optimized to minimize the negative effect of aminoacid replacements originated in mutational events [17,18]. In consequence, the application of modern algebraic tools involving finite fields and group theory, as applied in algebraic coding theory (for the design of error-correcting codes for the reliable transmission of information across noisy channels from human’s engineered communication systems) is forthright [16,19]. That is, the encoding of physicochemical or/and biophysical relationships between DNA bases and between codons is a mathematical tool to discover new knowledge about the transmission of genetic information, in analogous way as coding theory has been applied to human’s engineered communication systems.

Obviously, depending on which physicochemical or biophysical relationship is under scrutiny, different encodings of the group operations can be defined on the sets of bases and codons, as shown in reference [20]. The meaning of the group operations has been subjects of the references where the corresponding groups have been reported [11,20–23]. For example, in the DNA double helix, nucleotide bases are paired following specific physicochemical relationships: 1) *the chemical type sets the main rule for a paring: a purine base is paired with a pyrimidine*, 2) *paired bases must have the same hydrogen-bonding capability*. These physicochemical relationships rule the DNA base pairing: G:::C (*three hydrogen bonds*) and A::T (*two hydrogen bonds*). In this scenario, the sum operation is defined in [23], over the ordered set of bases 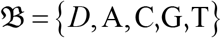, in such a way that the DNA complementary bases are also complementary algebraic elements.

#### Pioneering works on the genetic code algebraic structure

Pioneering works were made in the 70s [15,24–26], just few years after Nirenberg won the Nobel Prize in Physiology or Medicine (in 1968) for his seminal work on the genetic code. Andrews and Boss proposed the cyclic groups of DNA bases, which is isomorphic to the Abelian group defined on the set of integers modulo 4, 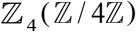 [15]. Their approach also considered the base representation with cyclic group of complex numbers. Further studies were focused on operational groups applied to transform bases and base-doublet into each other. Dankworth and Neubert (1975) proposed the Klein-4-group structure (*K*) of doublet-exchange operators and applied the direct product *K*×*K* to study the symmetries of genetic-code doublets [25]. The four dimensional hypercube structure of the genetic-code doublets (*K*×*K* group) was later studied by Bergman and Jungck [26].

Efforts to study the genetic code symmetries and symmetry breaking to gain understanding on genetic code degeneracies were made with the application of *group representation theory* to study the origin and evolution of the genetic code by Honors and Hornos and [27–29], Antoneli and Forger [30], and extended to Lie superalgebras by Forger and Sachse [31]. However, these efforts on the application of group representation theory are heavily relying on physical interpretations that made hard further applications on concrete computational biology and on bioinformatic applications. Here, it is important to recall that the *representation* of DNA bases, codons, genes, and genomic sequences as elements from algebraic structures must not be confused with the term *group representation* typically used in algebra referring to the theory of representations of algebraic structures or, particularly, the *group representation theory*. Nevertheless, once a group structure has been defined, for example, in the set of codons, a further application of the group representation theory can be developed.

In the current study, we aim to show that all possible genomic regions and, consequently, whole chromosomes can be described by way of finite Abelian groups which can be split into the direct sum of homocyclic 2-groups and 5-groups defined on the genetic code. Concepts and basic applications are introduced step by step, sometimes with self-evident statements for a reader familiar with molecular biology. However, it will be shown that the algebraic modeling is addressed to unveil more complex relationships between molecular evolutionary process and the genomic architecture than those eyes-visible relationships. This goal will be evidenced on section 3.2. Our algebraic model approach is intended to set the theoretical basis for further studies addressed to unveil and to understand the rules on how genomes are built. Concrete examples and an implementation in a R package are provided to pave the way for future computational and bioinformatic applications. A graphical summary of the modeling of DNA genomic regions proposed here is shown in Fig 1.

**Fig 1.**
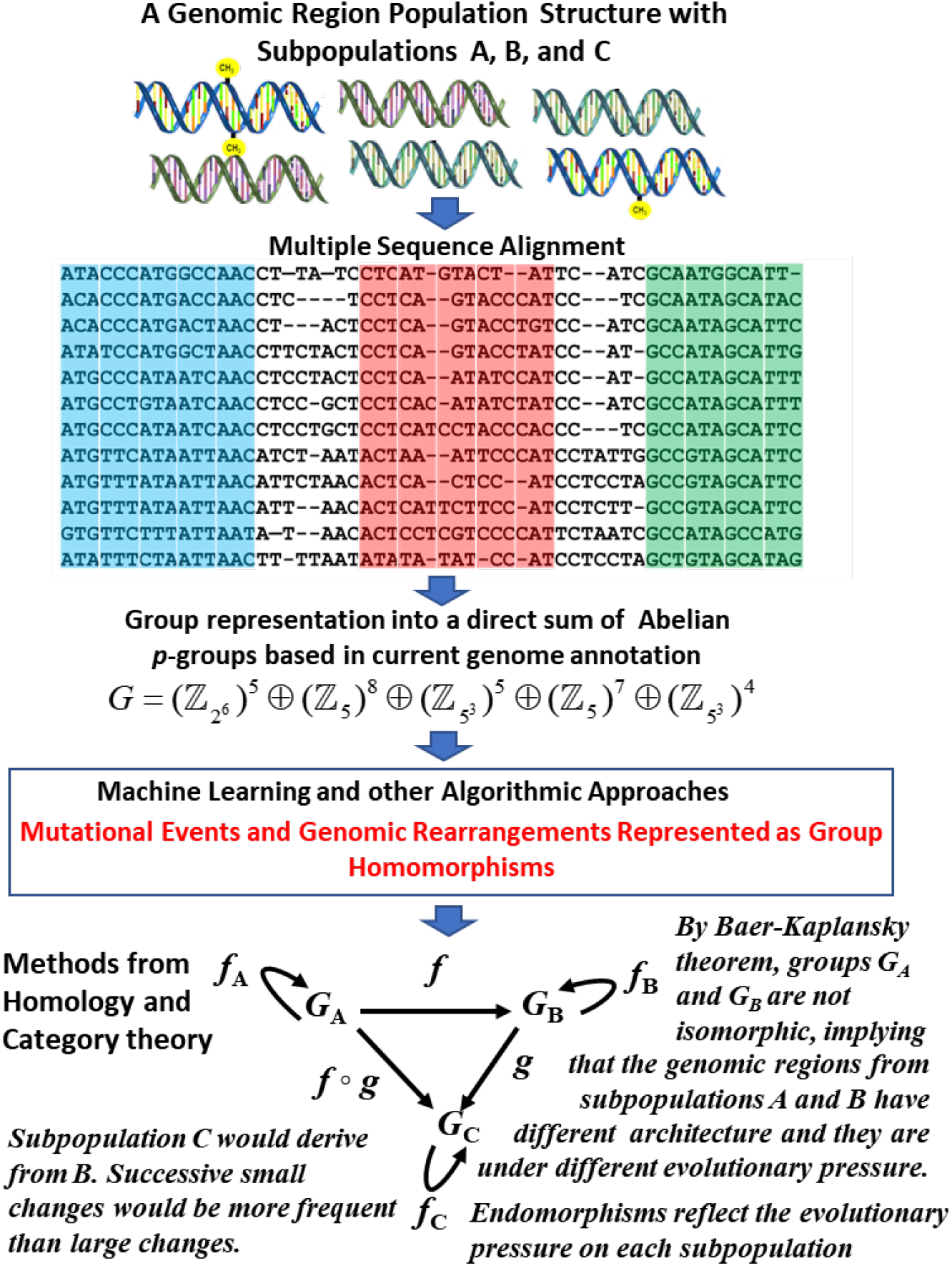
Graphical of the summary showing the bioinformatic and analytical steps followed in the algebraic modeling proposed in current work.

## 2 Materials and Methods

### 2.1 Preceding models applied in the current work

Of particular interest are the Abelian *p*-groups defined on the set of DNA bases 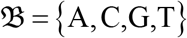 and on the set of 64 codons 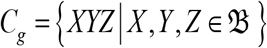, which are applied to modeling the physicochemical relationships between DNA bases in the codons [11,21]. Herein, for application purposes in computational biology and bioinformatics addressed to the study of the genome architecture, we focused our study on Abelian *p*-groups defined on 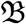 and on *C_g_* isomorphic to the groups 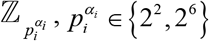, and on 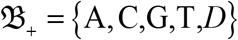 and 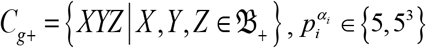, as presented in references [11,20–23].

Setting different physicochemical restrictions on the definition of groups operations leads to the 24 possible algebraic representations of the genetic code [20], i. e., 24 ways to encode the observed relationships between DNA bases and between codons. In particular, the Abelian *p*-group representations on the set 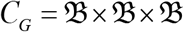 and 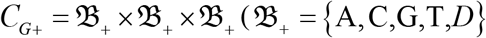, where *D* stands for an alternative base, see below) are isomorphic to Abelian groups defined on 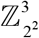 and 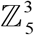, respectively. These group structures lead to 24 (isomorphic) geometrical representations of the genetic code as cubes inserted in three-dimensional space [11,20,22,23] (Fig 2 and SI Figs 1 and 3).

**Fig 2.**
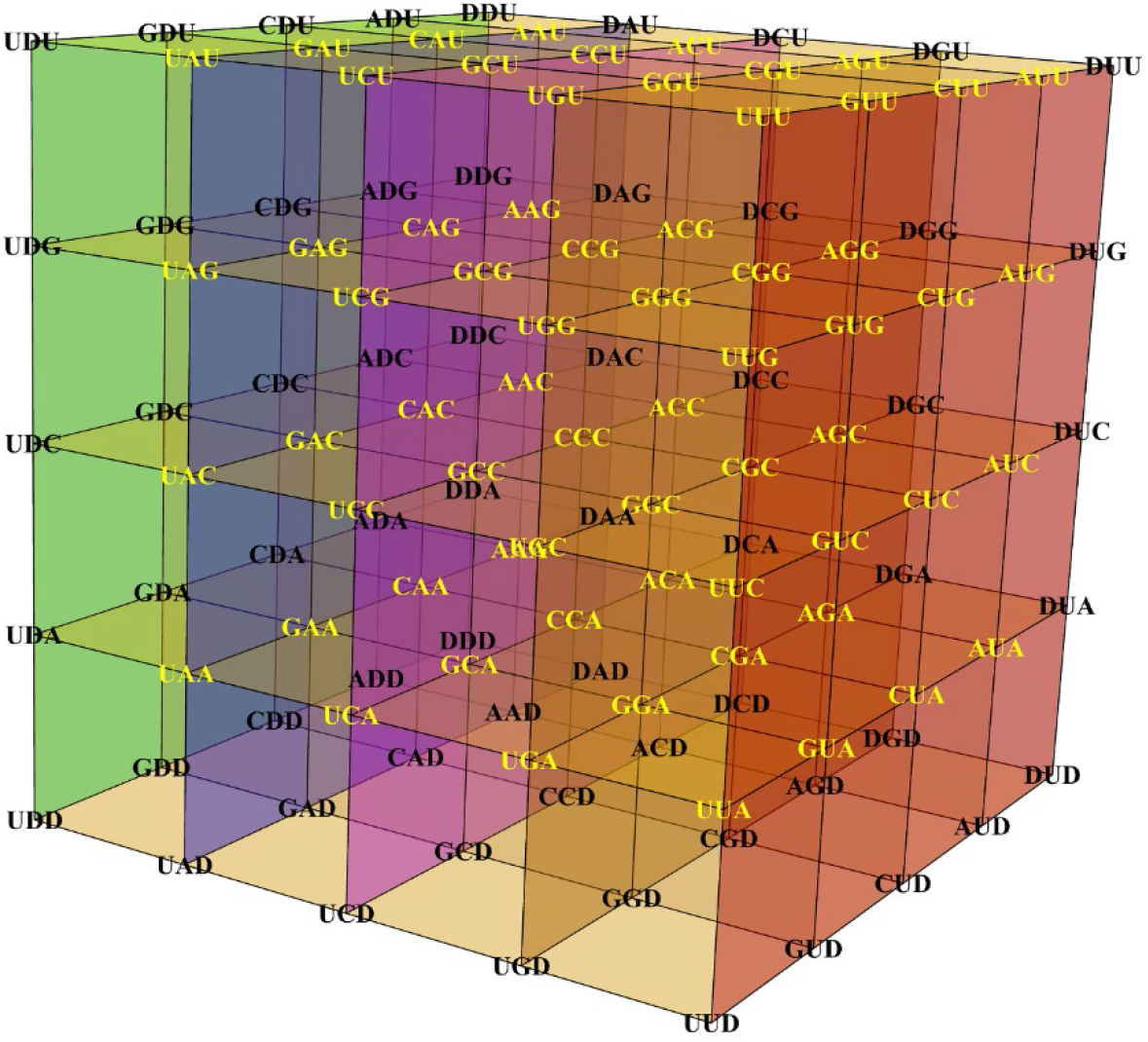
Geometrical representation of the genetic code as a cube inserted in three-dimensional space. The 2-group and 5-group representation defined on the sets 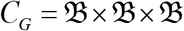and 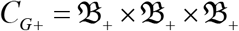 isomorphic to the groups defined on 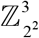 and 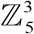, respectively, lead to the geometrical representations of the genetic code as a cube inserted in three-dimensional space. The cube corresponding to the base-triplets with coordinates on 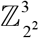 (yellow codons) is inserted in the cube with codon coordinates on 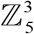. The extended base-triplets including the alternative base D (in black) are located on the cartesian coordinate planes. Codons encoding for amino acids with similar physicochemical properties are located on the same vertical plane (for more details on the cube description see also SI Fig 1 and reference [11,20,22,23]).

As shown in reference [11], a group structure isomorphic to the symmetric group of degree four *S*_4_ (preserving the group operations previously defined on the codon set) can be defined on the set the 24 genetic-code algebraic representations or in the set 24 cubes. Since the definition of a sum operation over the base set is equivalent to define an order on it, *cubes are named according to the base order on them*. For example, the cube shown in Fig 2 is denoted as ACGT, which correspond to the group operation defined on the ordered set 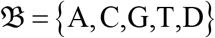 (the ‘dual’ cube TGCA is shown in SI Fig 2 [11]). Simulation of the evolutionary mutational process with the application of genetic algorithms indicates that fixed mutational events found in different protein populations are *very restrictive* in the sense that the optimal evolutionary codon distances are reached for specific models of genetic-code cube or for specific combination of genetic-code cube models [11]. In the present work, it will be shown that codon mutational events represented in terms of automorphisms can be also restrictive for specific genetic-code cube models (section 3.1).

All the Abelian *p*-group included in the current work are oriented to the study of the mutational process [11,20–23]. That is, since we are interested in those structures that permit the analysis and quantitative description of the mutational process in organismal populations, where mutational event can be represented by means of endomorphisms, automorphisms, and translations on a group. In the present work, we do not include algebraic structures designed to study the origin and evolution of the genetic code [11,21]. The genetic code is taken as currently is, without over-impose any evolutionary hypothesis on it.

A general model also consider Abelian 5-groups that includes a dummy variable (denoted by letter D), which extends the DNA alphabet to five letters. The usefulness of including a fifth base in the evolutionary analysis was shown in reference [23], where two evolutionary models, an algebraic and a stationary Markov (process) models, were applied to phylogenetic analysis reaching (both models) greater discriminatory power than the (now) classical Tamura-Neil evolutionary (Markov) model based on four DNA alphabet [32]. Depending on the concrete application, letter “D” will take a different value. The possible values in the context of the present modeling are: 1) the gap symbol “-”, which stands for insertion deletion/mutations in the multiple sequence alignment (MSA) of DNA sequences, 2) alternative wobble base pairing (e.g., bases such as: inosine (in eukaryotes), agmatine (in archaea), and lysidine (in bacteria) [17,21,22]), and 3) 5-methylcytosine (C^m^) and N-6-methyladenine (A^m^) when intended for epigenetic studies.

A concrete application of the extended genetic-code cubes over the Galois field *GF*(5) to the simulation of the mutational process proposed in reference [11] would be particularly relevant to predict immunoescape epitope variants originated in populations of pathogenic microorganisms and viruses. In addition, examples provided (here) on the application of the algebraic model to DNA methylation (on 5-methylcytosine and on N-6-methyladenine) suggest its importance for epigenetic studies. The analysis of the fixed mutational events on genes populations revealed that the mutational process can be described by automorphisms on different cubes or sets of cubes [11]. The best genetic-code cubes describing the mutational process on a given gene population are selected with the application of an optimization algorithm (evolutionary (genetic) algorithms) using multiple sequence alignment as raw data [11].

It is worthy to notice that, for all mentioned Abelian *p*-groups, the calculus can be accomplished as symbolic computation on the set of DNA bases or on the set of codons (see e.g., [21]). However, for practical purposes, we take advantage of the group isomorphisms. That is, after define group structures on the sets of bases and codons, for the sake of straightforward computation it is convenient to take advantage of the group isomorphisms with the Abelian *p*-groups like: 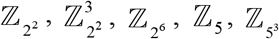 and 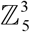, which will be used in our study instead of the original groups defined on the sets of bases and codons (base-triplets). An introductory summary on the mentioned algebraic structure defined on the set of codons is provided as supporting information in S1.

In the context of genetic-code algebraic structures, by the term “*representation*” of DNA bases, codons, genes, and genomic sequences as elements from algebraic structures, we understand the symbolic representation of the mentioned biomolecules and the physicochemical relationships between them by means of group operations defined on the given set of biomolecules.

### 2.2 Aligned DNA sequences and data sets

All the DNA sequence alignments and data sets used in this work are available within the R package *GenomAutomorphism* (version 1.0.0) [33]. In addition, the pairwise sequence alignments of SARS coronaviruses used the analyses shown in Fig 8**a** and **b** are also available at GitHub in: https://github.com/genomaths/seqalignments/tree/master/COVID-19. The multiple sequence alignment (MSA) of primate somatic cytochrome c and data description are available on GitHub at: https://github.com/genomaths/seqalignments/tree/master/CYCS. This MSA includes DNA protein-coding sequences from: human, gorilla, silvery gibbon, white cheeked gibbon, Francois langur, olive baboon, golden monkey, rhesus monkeys, gelada baboon, and orangutan. The MSA of primate BRCA1 (transcript variant 4) DNA repair gene used to compute the automorphism shown Fig 8**d** is available on GitHub at https://github.com/genomaths/seqalignments/tree/master/BRCA1. The MSA, coordinates and R script to create the sequence-logo from Fig 4 are given in the Supporting Information.

**Fig 3.**
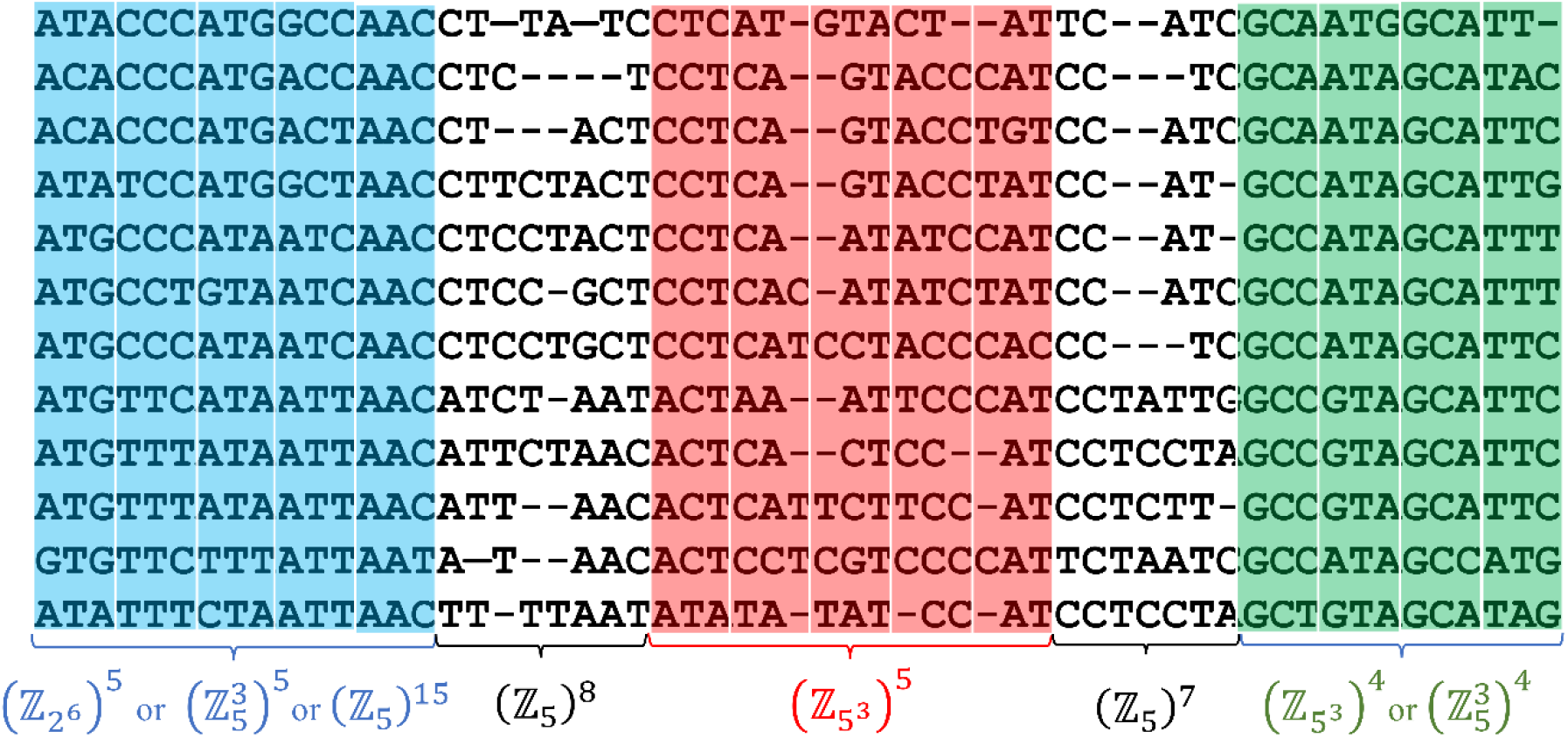
An illustration of a typical DNA multiple sequence alignment (MSA) including segments of protein-coding regions. A MSA would include the presence of substitution, insertion, and deletion mutations (*indel* mutations). The aligned sequences can be grouped into blocks, which can be algebraically represented by Abelian groups. A homocyclic group covering a MSA block corresponds to a sub-classification of the protein-coding region into subregions and, consequently, leading to a more accurate molecular taxonomy of species. In protein-coding regions cyclic groups 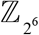 and 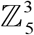 are appropriated to study exon regions, while 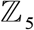 for non-coding intron regions. As shown in section 1.4, the group representation leads us the analysis of the more frequent mutational events (represented as endomorphisms and translations) observable in genes from organismal populations.

**Fig 4.**
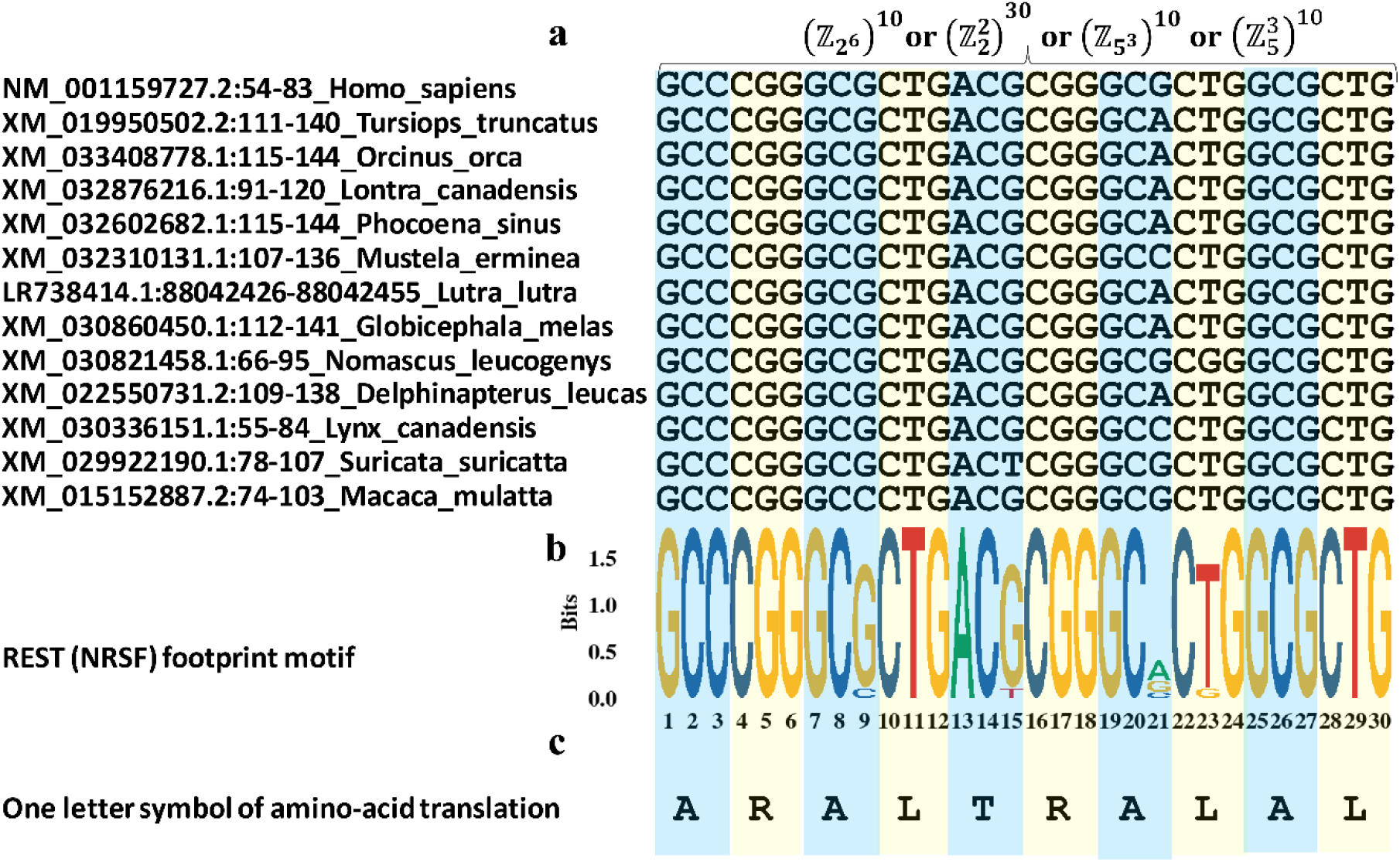
The DNA sequence motifs targeted by transcription factors usually integrate genomic building block across several mammal species. **a**, DNA sequence alignment of the protein-coding sequences from phospholipase B domain containing-2 (PLBD2) carrying the footprint sequence motif recognized (targeted) by the Silencing Transcription factor (REST), also known as Neuron-Restrictive Silencer Factor (NRSF) REST (NRSF). **b**, Sequence logo of the footprint motif recognized REST (NRSF) on the exons. **c**, Translation of the codon sequences using the one-letter symbol of the aminoacids.

**Fig 5.**
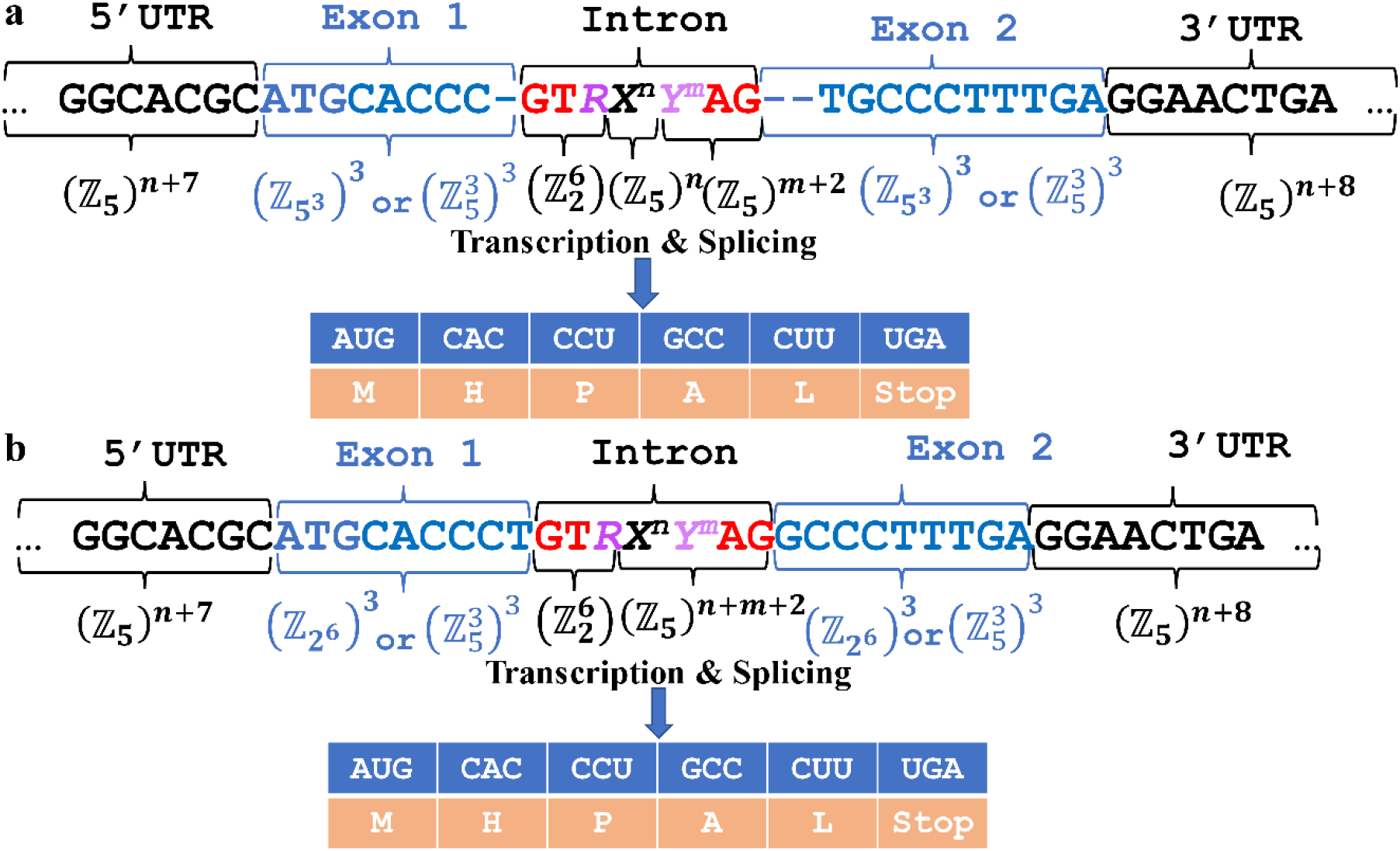
Two different protein-coding (gene) models can lead to the same Abelian group representation and the same protein sequence. A dummy intron was drawn carrying the typical sequence motif targeted by the spliceosome donor (GU*R*) and acceptor (*Y^m^* AG) sites, where *R* ∈ {A, G} (purines) and *Y* ∈{C, U}, *X* stands for any base, and *n* and *m* indicate the number of bases present in the corresponding sub-sequences (pyrimidines). **a**, A gene model based on a *dummy* consensus sequence where gaps representing base D from the extended genetic code were added to preserve the coding frame, which naturally is restored by splicing soon after transcription. **b**, A gene model where both exons, 1 and 2, carries a complete set of three codons (base-triplets). Both gene models, from panels **a** and **b**, share a common group representation as direct sum of Abelian 5-groups.

**Fig 6.**
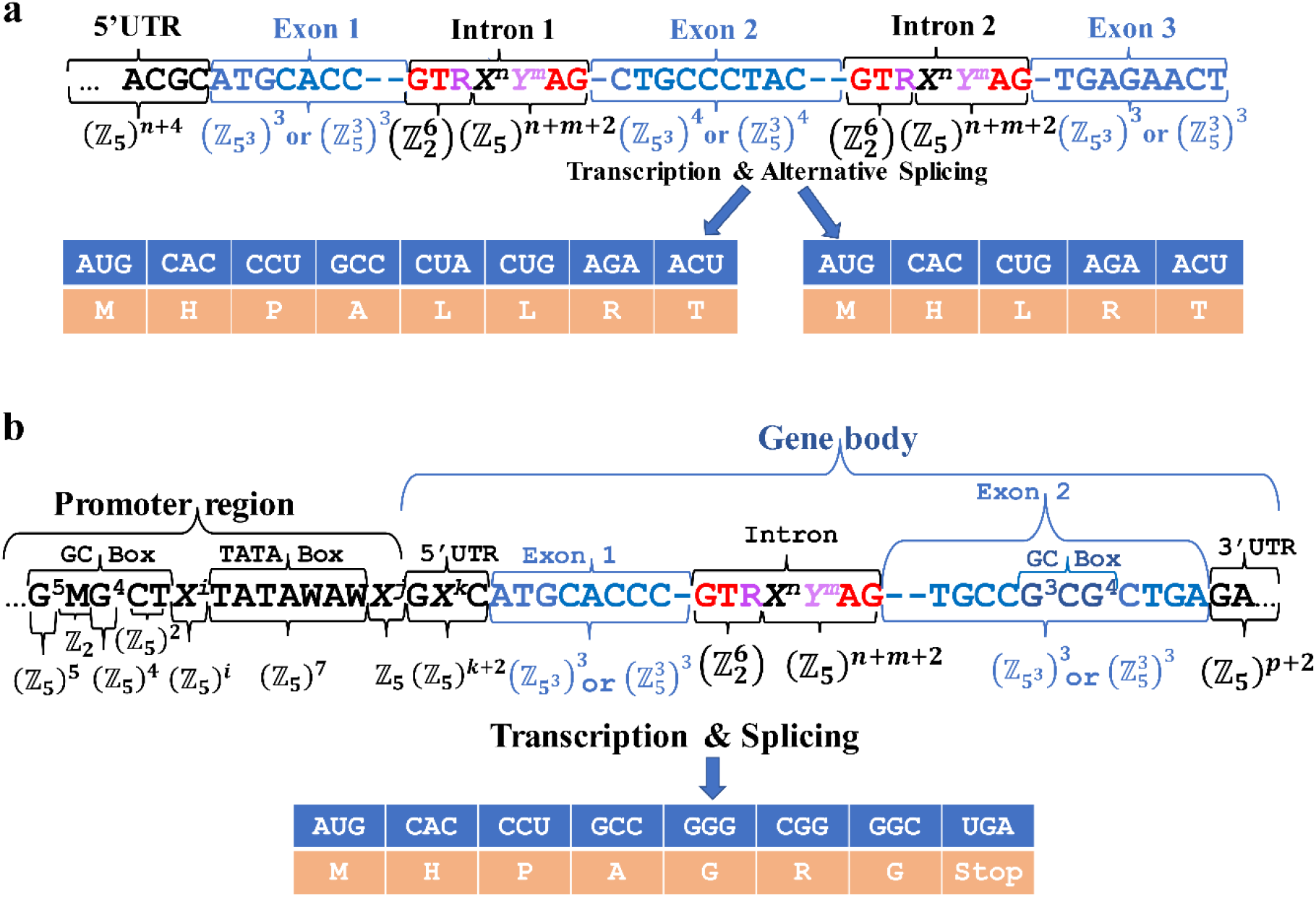
The Abelian group representation of a given genome takes advantage of our current knowledge on its annotation. **a,** the alternative splicing specified for an annotated gene model does not alter the Abelian group representation and only would add information for the decomposition of the existing cyclic groups into subgroups. **b**, a more complex gene model including detailed information on the promoter regions. A GC box (G5MG4CU) motif is located upstream of a TATA box (TATAWAW) motif in the promoter region. The GC box is commonly the binding site for Zinc finger proteins, particularly, Sp1 transcription factors. A putative GC box was included in exon 2, which is an atypical scenario, but it can be found, e.g., in the second exon from the gene encoding for sphingosine kinase 1 (SPHK1), transcript variant 2 (NM_182965, CCDS11744.1). In this group representation, the spliceosome donor GT*R* can be represented by the elements from a quotient group (see main text).

**Fig 7.**
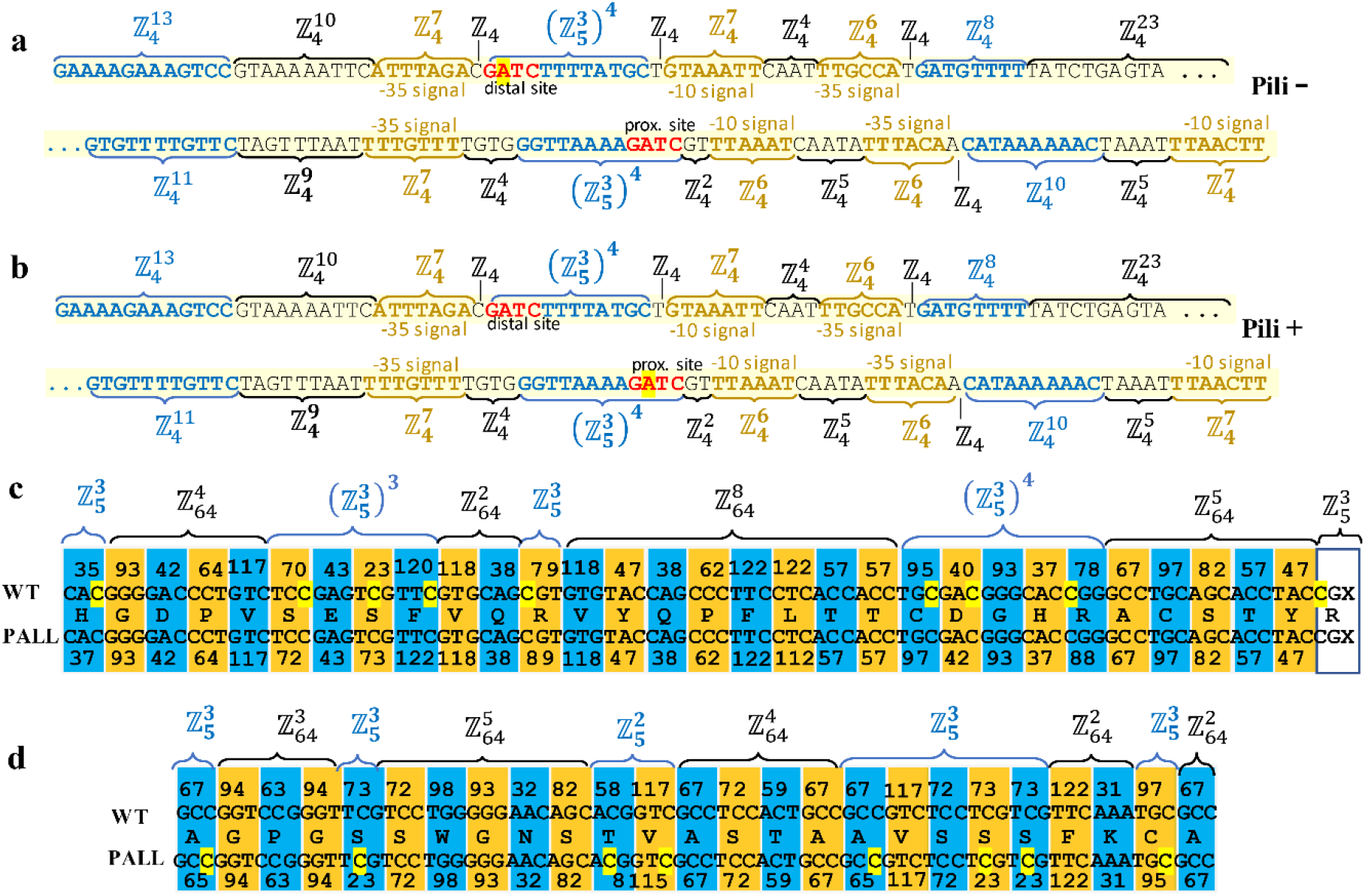
Vector representation of differentially methylated gene regions. **a** and **b**, regulation of pyelonephritis-associated pilus (pap) expression by DNA methylation on the Escherichia coli operon (locus X14471). **c** and **d**, exons regions from genes EGEL7 and P2RY1 from patients with pediatric acute lymphoblastic leukemia (PALL). In panel **a**, two 5’-GATC-3’ DNA adenine methyltransferase (Dam) methylation sites in the middle of each set of the leucine-responsive regulatory protein (Lrp) binding sites (in blue). In the inactive state, panel **b**, a Lrp octamer is bound to the three proximal Lrp 3’ sites, while the GATC^dist^ site in Lrp site 5 is fully methylated, and the system remains in phase OFF (Pili -) regarding pilus expression. In the active state, the adenine from the GATC^prox^ is methylated permitting to bend the DNA to recruit CRP to activate transcription of papBA genes (Pili +). Pap pili are multisubunit fibers essential for the attachment of uropathogenic Escherichia coli to the kidney (see [45]). In panel **c**, a segment of exon-6 from gene EGFL7 located at chromosome 9: 139,563,008-139,563,124 is shown. On average, this gene is hypo-methylated in the control group with respect to PALL group. **d**. Segment of exon-1 from gene P2RY1. Methylated cytosines are highlighted in yellow background. In PALL patients, gene EGEL7 mostly hypomethylated and gene P2RY1 mostly hypermethylated in respect to healthy individuals (WT). The encoded aminoacid sequence is given using the one letter symbols. Both genes, EGEL7 and P2RY1, were identified in the top ranked list of differentially methylated genes integrating clusters of hubs in the protein-protein interaction networks from PALL reported in reference [46]. The integer number at the top and bottom of panel **c** and **d** stand for the codon coordinates in 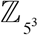 (see SI Table 1).

**Fig 8.**
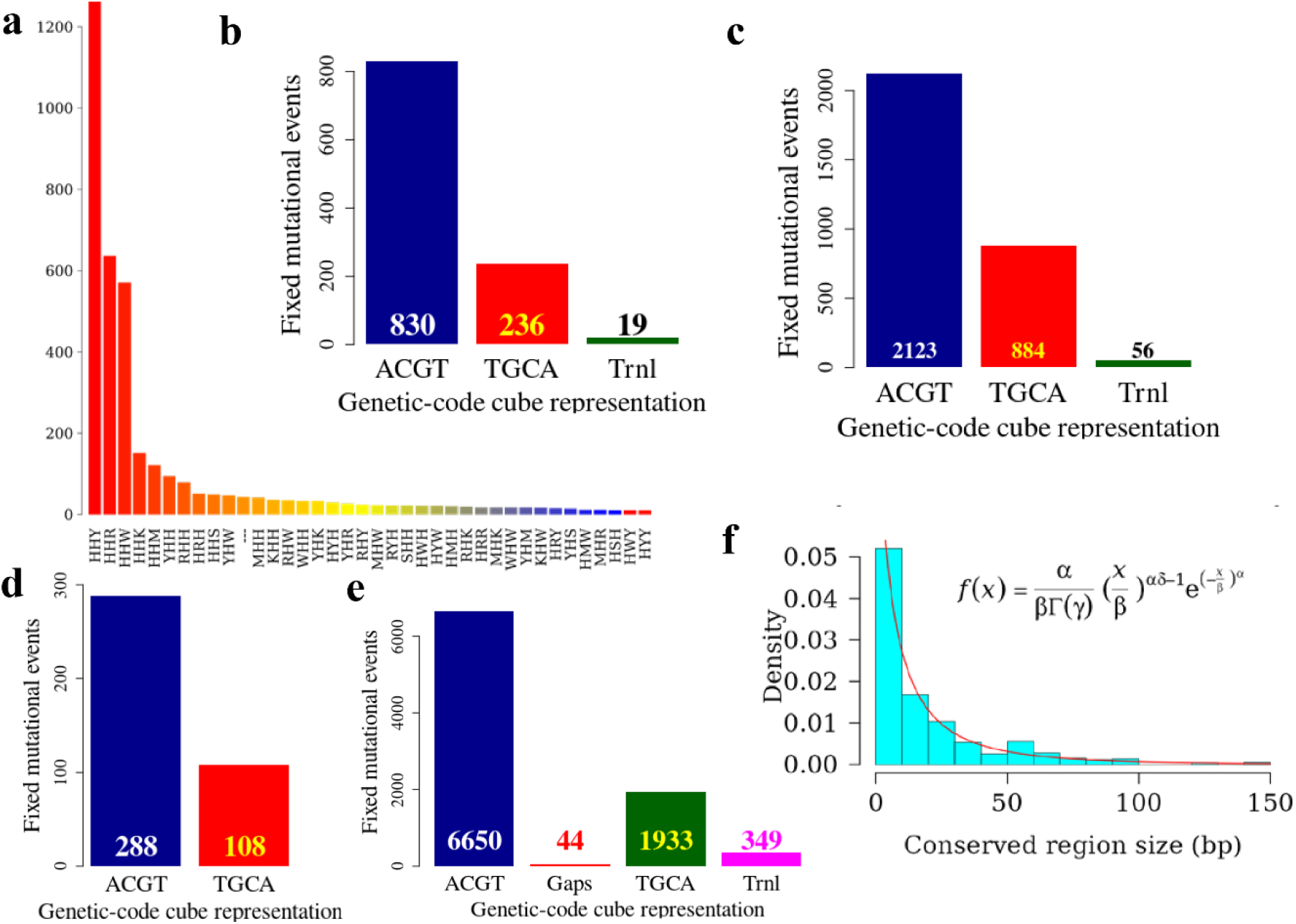
Analysis of mutational events in terms of automorphisms on DNA protein-coding regions represented as homocyclic groups on 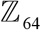. In the Abelian group defined on 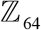, automorphisms are described as functions *f*(*x*) = *k x* mod 64, where *k* and *x* are elements from the set of integers modulo 64. **a**, Frequency of mutational events (automorphisms) according to their mutation type. That is, every single base mutational event across the MSA was classified according IUPAC nomenclature [51]: 1) According to the number of hydrogen bonds (on DNA/RNA double helix): strong S={C, G} (three hydrogen bonds) and weak W={A, U} (two hydrogen bonds). According to the chemical type: purines R= {A, G} and pyrimidines Y= {C, U}. 3). According to the presence of amino or keto groups on the base rings: amino M= {C, A} and keto K= {G, T}. Constant (hold) base positions were labeled with letter H. So, codon positions labeled as HKH means that the first and third bases remains constant and mutational events between bases G and T were found in the second codon position. **b** and **c**, Bar plots showing the frequency of automorphisms found on the *group of dual cubes* (see [11]): ACGT – TGCA and CATG – GTAC on 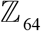 between SARS coronavirus GZ02 and bat SARS-like coronaviruses: **a**, isolate Rs7327 (GenBank: KY417151.1, protein-coding regions) and **c**, isolate bat-SL-CoVZC45 (GenBank: MG772933.1:265-1345513455-21542, nonstructural polyprotein). **d**, frequency of automorphisms between human somatic cytochrome c and other nine primates (monkeys). **e**, frequency of automorphisms between human BRCA1 DNA repair gene and other seven primates (see Material and Method section). **f**, Distribution of the conserved COVID-19 genomic regions according to their size. The graphics result from the analysis SARS coronavirus GZ02 versus the two mentioned bat strains. The best fitted probability distribution turned out to be the generalized gamma distribution.

### 2.3 Software applied for the mathematical and statistical analyses

Results shown in Fig 8 and Fig 9 were obtained applying the *GenomAutomorphism* R package [33] (version 1.0.0), which is available at Bioconductor (the open source software for Bioinformatics, version: 3.16) and, also, in GitHub at: https://github.com/genomaths/GenomAutomorphism. The whole R script pipeline applied in the estimation of automorphisms (Fig 8) and decision tree (Fig 9) are available as tutorials (vignettes) at the *GenomAutomorphism* website: https://github.com/genomaths/GenomAutomorphismm.

**Fig 9.**
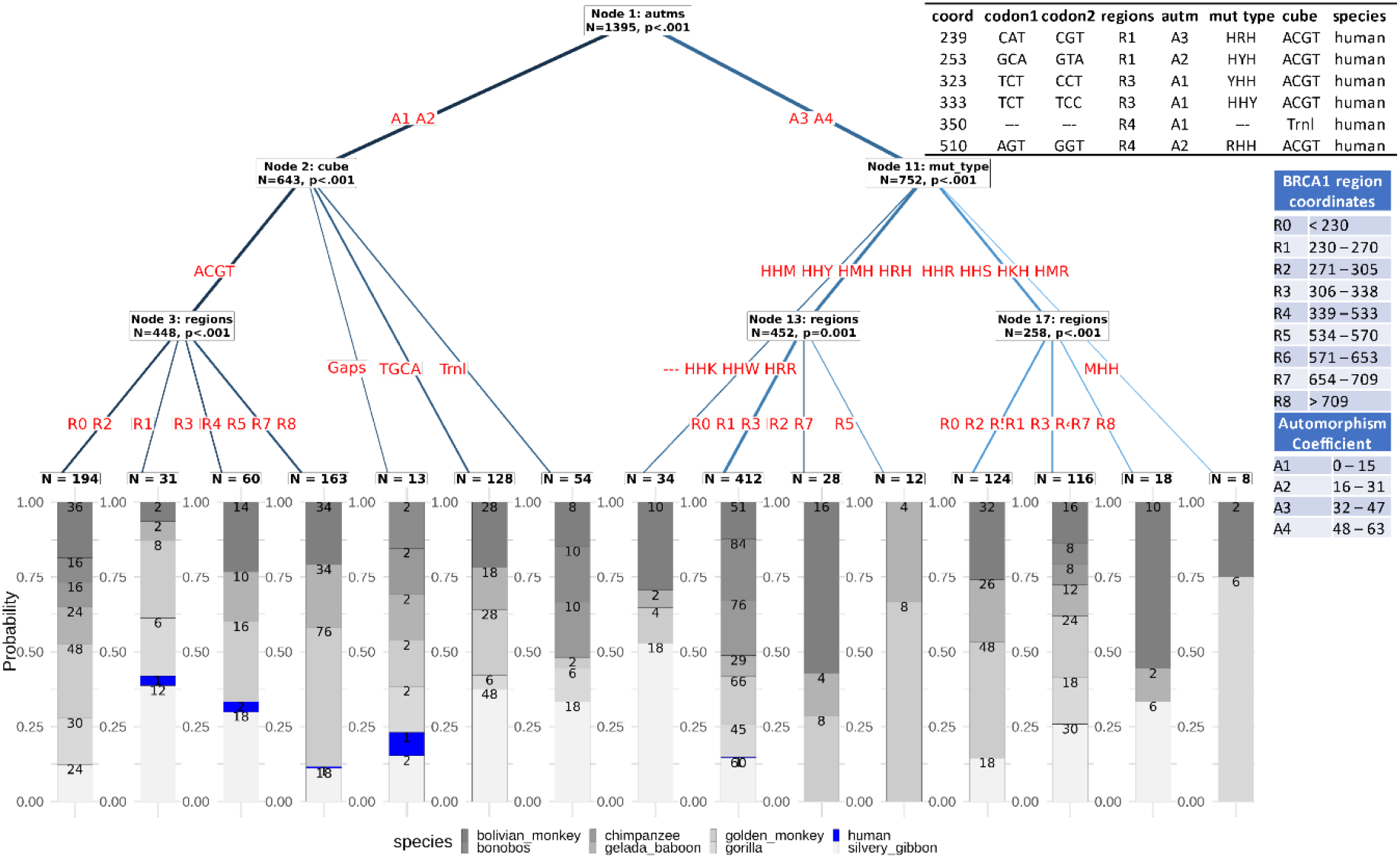
Decision tree based on automorphisms estimated on primate BRCA1 genes. Symbols R0 to R8 denote the protein regions as given in UniProt plus inter regions segments (see https://www.uniprot.org/uniprot/P38398#family_and_domains). Only regions experiencing fixed mutational events are included in the analysis. The range of automorphism coefficients *k*(*f*(*x*) = *k x* mod 64) are denoted after the isomorphism between the genetic-code cyclic group defined in the set of codons and the Abelian group defined on 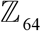. For the sake of graphic comprehension, the coordinates of human-to-human mutations were added. Every branch (path) from the top to the leaf node is equivalent to a stochastic-determinist logical rule defining the automorphism preference for each protein region in the subset of analyzed primate BRCA1 genes. For example, with high probability the rule: “(A4 ˅ (A3 ˄ ¬ R1)) → ¬ human” is true (see Supporting Information).

The estimation of the best fitted probability distribution shown in Fig 8**f** was accomplished with R package *usefr* available at GitHub: https://github.com/genomaths/usefr, and the goodness-of-fit tests are reported in the mentioned tutorials. The genetic-code cube shown in Fig 2 was obtained from the Wolfram Mathematica Notebook: *Introduction to* 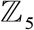-*Genetic-Code vector space*, free available at https://github.com/genomaths/GenomeAlgebra_SymmetricGroup. All the examples of modular matrix operations included in the manuscript can be reproduced following a tutorial available at *GenomAutomorphism* website.

### 2.4 Theoretical Model

According to the fundamental theorem of Abelian finite groups (FTAG) [34,35], any finite Abelian group can be decomposed into a direct sum of homocyclic *p*-groups [34], i.e., a group in which the order of every element is a power of a primer number *p*. Herein, it will be shown that, in a general scenario, genomic regions and, consequently, whole genome populations from any species or close related species, can be algebraically represented as a direct sum of Abelian homocyclic groups or more specifically Abelian *p*-groups of *prime-power order*. The multiple sequence alignments (MSA) of a given genomic region of *N* base-pair (bp) length can be represented as the direct sum:

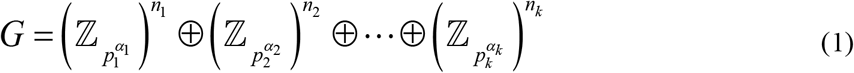

Where 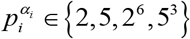, *n_i_* stands for the number of cyclic groups 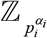 integrating the homocyclic group 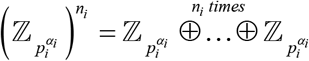. Here, we assume the usual definition of direct sum of groups [35]. For 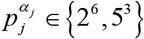 the cyclic group 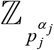 will cover three bases, otherwise only one base (see examples below). Considering such groups (not necessarily in the order given in Eq. 1) we have: *N* = *n*_1_ +… + *n_j_* + *n*_*j*+1_ +… + *n_j+m_* +… + *n_k_*. Throughout the exposition of the theory and examples given in the next sections, it will be obvious that the group representations can be extended, starting from small genomic regions till cover whole chromosomes and, consequently, the whole genome, i.e., the set of all chromosomes.

Let *B_i_* (*i* ∈ *I* = {1,…,*n*}) be a family of subgroups of *G*, subject to the following two conditions:

1. ∑*B_i_* = *G*. That is, *B_i_* together generate *G*.
2. For every *i* ∈ *I* and *i* ≠ *j*: *B_i_* ⋂ ∑*B_j_* = 0.

Then, it is said that *G* is the direct sum of its subgroups *B_i_*, which formally is expressed by the expression: 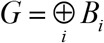 or *G* = *B*_1_ ⊕… ⊕ *B_n_*.

Genomic DNA sequences from superior organisms are integrated by intergenic regions and gene regions. The former are the larger regions, while the later includes the protein-coding regions as subsets. The MSA of DNA and protein-coding sequences reveal allocations of the nucleotide bases and aminoacids into stretched of *strings*. The alignment of these stretched would indicate the presence of substitutions, insertions, and deletion (*indel*) mutations. As a result, the alignment of homolog genomic regions or whole chromosome DNA sequences from several individuals from the same or close-related species can be split into well-defined subregions or domains, and each one of them can be represented as homocyclic Abelian groups, i.e., as the direct sum of cyclic group of the same *prime-power* order (Fig 3). As a result, each DNA sequence is represented as a *N*-dimensional vector with numerical coordinates representing bases and codons.

An intuitive mathematical representation of a MSA is implicit in Fig 3, with the following observations:

a. Bases or codons can be represented as elements of an Abelian group defined on the set of bases or on the set of codons. In the second block (including gap symbol ‘-’) each base from each sequence is represented as an element from the Abelian group defined on the set {A, C, G, T, D }, where D = ‘–’, which is isomorphic to the Abelian *p*-group defined on the set 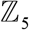
b. The extended base triplets (including gaps symbol ‘-’) from each sequence in the third aligned block are represented as elements from the Abelian *p*-group defined on the set of extended base-triplets (125 element, see SI Table 1) which is isomorphic to the Abelian group defined on the set 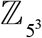, and so on.
c. Every DNA sequence from the MSA and every subsequence on it can be represented as a *numerical vector* with element coordinates defined in an Abelian group. For practical computational purposes we take advantage of the group isomorphism to work with numerical representations of DNA bases and codons. For example, codons from the first aligned block (in blue) can be represented as elements from an Abelian group defined on the set of codons, which can be isomorphic to 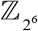 or to 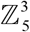. That is, since 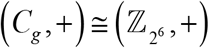, the first five codons {ATA, CCC, ATG, GCC, AAC} ∈ *C_g_* from the first DNA sequence from Fig 3, can be represented by the vector of integers: {48,21,50,25,1} where each coordinate is an element from group 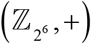 (see Table 1 from reference [21]).
d. Any MSA can be algebraically represented as a symbolic composition of Abelian groups each one of them is isomorphic to an Abelian group of integers module *n*. Such a composition can be algebraically represented as a direct sum of homocyclic Abelian *p*-groups. For example, the MSA from Fig 3 can be represented by the direct sum of five homocyclic Abelian *p*-groups:

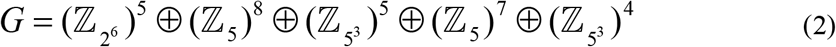 Where the length of each region determines the number of cyclic *p*-groups in the corresponding homocyclic Abelian *p*-group 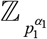 representing each region. For example, in Eq. 2 we have the homocyclic group: 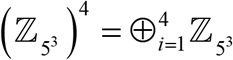, which is a direct sum of 4 cyclic 5-groups 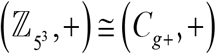. Since group *G* is the direct sum of homocyclic Abelian *p*-groups of different prime-order, we shall say that *G* is a heterocyclic group.

In more specific scenario, the MSA from Fig 3 can be represented by only one homocyclic

Abelian 5-group:

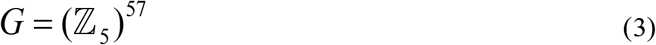

But this representation ignores the local variability detected by the MSA algorithm. Hence, preserving the highlighted features, the MSA can be represented as the direct sum of homocyclic Abelian 5-groups:

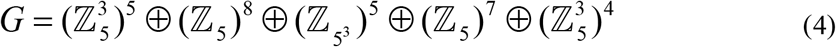

Although the above *direct sums* of Abelian *p*-groups provides a useful compact representation of a MSA, for application purposes to genomics, we would also consider to use the concept of direct product (*cartesian sum or complete direct sums*) [35]. Next, let *S* be a set of Abelian cyclic groups identified in a MSA *M* of length *N* (i.e., every DNA sequence from *M* has *N* bases). Let *ℓ_i_* the number of bases or triples of bases covered on *M* by group *S_i_* ∈ *S* where ∑_*i*_*ℓ_i_* = *N*. Hence, each DNA sequence on the *M* can be represented by a cartesian product (*b*_1_,…, *b_n_*) where *b_i_* ∈ *S_i_* (*i* = 1,…, *n*) and *n* = |*S*|. Let 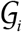 be a group defined on the set of all elements (0,…, 0, *b_i_*, 0,… 0) where *b_i_* ∈ *S_i_* stands on the *i^th^* place and 0 everywhere else. It is clear that 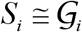. In this context, the set of all vectors (*b*_1_,…, *b_n_*) with equality and addition of vectors defined coordinate-wise becomes a group 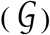 named direct product (cartesian sum) of groups 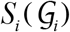, i.e.:

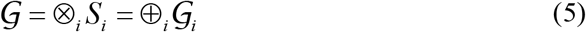

An illustration of the cartesian sum application was given above in observation a).

## 3 Results

Results essentially comprise an application of the fundamental theorem of Abelian finite groups [34,35]. By this theorem, every finite Abelian group *G* is isomorphic to a direct sum of cyclic groups of prime-power order of the form:

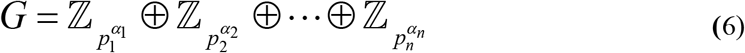

Or (in short) 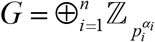, where the *p_i_*’s are primes (not necessarily distinct), 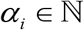 and 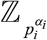 is the group of integer module 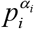. The Abelian group representation of the MSA from Fig 3 given by Eq. 2 correspond to a heterocyclic group that split into a direct sum of homocyclic Abelian 2-groups and 5-groups, each one of them split into the direct sum of cyclic *p*-groups with same order; while in Eqs. 3 and 4, the Abelian group *G* is decomposed into a direct sum of homocyclic Abelian 5-groups [34,35].

Notice that for a large enough genomic region of fixed length *N* we can build a *manifold of (a set of various) heterocyclic groups S_i_*, where each one of them can have different decomposition into *p*-groups. The set *S* of all possible Abelian *p*-group representations *S_i_* of a large genomic region of fixed length (having numerous different parts, elements, features, forms, etc.) that split into the direct sum of several heterocyclic groups 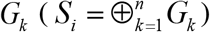 shall be called a *heterocyclic-group manifold*. So, each genomic region can be characterized by means of their corresponding *heterocyclic-group manifold*.

### 3.1 Examples of genomic regions group representations

A group representation is particularly interesting for the analysis of DNA sequence motifs, which typically are highly conserved across the species. As suggested in Fig 3 and 4, there are subregions of DNA or protein sequences where there are few or not gaps introduced and mostly substitution mutations are found. Such subregions conform blocks that can cover complete DNA sequence motifs targeted by DNA biding proteins like transcription factors (TFs, Fig 4), which are identifiable applying bioinformatic algorithms like BLAST [36].

The case of group representation on a TF binding motif is exemplified in Fig 4, where an exon region from the enzyme *phospholipase B domain containing-2* (PLBD2) simultaneously encodes information for several aminoacids and carries the footprint to be targeted by the transcription factor REST. Herein, the case of double encoding called our attention, where the DNA sequence simultaneously encodes the information for transcription enhancer target motif and for a codon sequence (base-triplets) encoding for aminoacids. These types of double-coding regions are also called *duons* [37–39].

Four group representations for this exon subregion are suggested in the top of the Fig 4 (panel **a**). However, the MSA’s sequence logo (panel **b**) suggests that this transcription factor binding-motif is a highly conserved codon sequence in mammals (with no indel mutations on it) and, in this case, the Abelian group 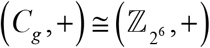 defined on the standard genetic code is the appropriated model to represent these motifs (Fig 4). The homocyclic group representation of conserved and biological relevant DNA sequence motifs, illustrated in Figs. 3 and 4, stablish the basis for the study of the molecular evolutionary process in the framework of group endomorphisms and automorphisms as suggested in [21,23] (section 1.4).

In Fig 5, two different protein-coding (gene) models from two different genome populations can lead to the same direct sum of Abelian *p*-groups and to the same final aminoacids sequence (protein).

The respective exon regions have different lengths and gaps (“-”, representing base D in the extended genetic code) were added to exons 1 and 2 (from panel **a**) to preserve the reading frame in the group representation (after transcription and splicing gaps are removed). Both gene models, from panel **a** and **b**, share a common direct sum of Abelian 2-groups and 5-groups: 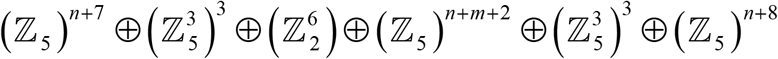. The analysis of theses gene models suggests that *DNA sequences sharing a common group representation as direct sum of Abelianp-groups would carry the same or similar, or close related biological information*. However, it does not imply that the architecture of these protein-coding regions is the same. The gene model in panel **b** permits the direct sum representation: 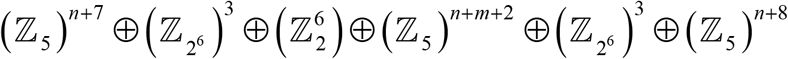, which is no possible for the gene model from panel **a**. That is, the *heterocyclic-group manifold* from the gene model in panel **a** is different from the one in panel **b**. The difference of group representation just captures the obvious fact that these gene models are different and, consequently, their gene architectures are different.

At this point we shall introduce the concept of *equivalent class of genomic region*. We shall say that two genomic regions belong to same *equivalent class of genomic region* if they hold the same heterocyclic-group manifold (and, consequently, they hold same architecture). Under this definition, the region architecture of the protein-coding regions from Fig 5**a** and **b** are not equivalent. The concept of *equivalent class of genomic region* is relevant for further applications of the group representation on the taxonomy study of organismal populations.

Taxonomy is the study of the scientific classification of biological organisms into groups based on shared characteristics. Mathematically, this is a way to split biological organisms into classes of equivalences. Numerical taxonomy is a well-established application of multivariate statistics on the analysis of plant germplasm banks. The group representations of genomic regions will lead to a higher accuracy in the taxonomy study of organismal populations.

No matter how complex a genomic region might be, it has an Abelian group representation (as described above). A further application of group theory would unveil more specific decomposition of small genomic regions into Abelian groups. For example, the set of base-triplets found in a typical sequence motif targeted by the spliceosome donor, GT*R* (Figs. 5 and 6), is in the vertical line GT*Z* (GU*Z*) of the vertical plane *X*T*Z* (*X*U*Z*) from the cube ACGU shown in Fig 2 (see also SI Fig 3).

Since purine bases (R: A and G) are the only accepted variants at the third codon position, it is convenient to model these base-triples with the group defined on the cube AGCU [11] (SI Fig 3). Next, following analogous reasoning as in [22], it turns out that the set of base-triplets GT*R* is a coset from the quotient group (*C_G_*, +)/*G*_AAG_, where here 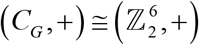 is the additive group from the genetic code Galois field *GF*(64) reported in reference [40] and *G*_AAG_ = ({AAA,AAG}, +) is a subgroup from the Klein four group defined on the set {AAA,AAG,AAC,AAU} (see operation table in the SI Table 2), i.e., GT*R* = GTA + *G*_AAG_ (SI Fig 3).

There exists strong evolutionary pressure on splicing donor site to keep the base-triplet GTR in the genetic-code cube vertical line GTZ (GUZ). As shown in the clinical report [41] mutational variants, located in different cube’s vertical lines (different cosets, SI Fig 3) GCZ and CTZ (CUZ), within intron 3 have led to four aberrant RNAs transcripts that causes rare X-chromosome-linked congenital deafness. As will be shown below (in section 3.1) the strong connection between DNA sequences and non-disrupting mutational events is mathematically (and accurately) modeled by the strong relationship between a group representation and the endomorphism ring on it.

An example considering changes on the gene-body reading frames as those observed in alternative splicing is shown in Fig 6. Gene-bodies with annotated alternative splicing can easily be represented by any of the groups 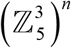 or 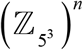 (Fig 6**a**). The splicing can include enhancer regions as well (Fig 6**b**) [42]. Enhancers are key regulator of differential gene expression programs.

As commented in the introduction, cytosine DNA methylation is implicitly included in extended base-triple group representation. Typically, the analysis of methylome data is addressed to identify methylation changes induced by, for example, environmental changes, lifestyles, age, or diseases. So, in this case the letter D stands for methylated adenine and cytosine (D = *C^m^*), since only epigenetic changes are evaluated.

Concrete examples of adenine in bacteria linked to the regulation of pyelonephritis-associated pilus (pap) expression by DNA methylation on the *Escherichia coli* operon (locus X14471) and cytosine methylation in two (humans) genes from patients with pediatric acute lymphoblastic leukemia (PALL) are presented in Fig 7. On protein-coding regions methylation changes can be analyzed on the homocyclic groups composed by the cyclic group 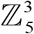 or 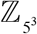 (Fig 7c and **d**). Notice that adenine methylation is found in humans as well and, usually, it plays a very specific regulatory role [43,44].

It is obvious that the MSA from a whole genome derives from the MSA of every genomic region, from populations from the same or closed related species. At this point, it is worthy to recall that there is not, for example, just one human genome or just one from any other species, but populations of human genomes and genomes populations from other species. Since a relatively small genomic region can be represented by the direct sum of Abelian homocyclic groups of prime-power order, then the whole genome population from individuals from the same or closed related species can be represented as an Abelian group, which will be, in turns, the direct sum of Abelian homocyclic groups of prime-power order. Hence, results lead us to the representation of genomic regions from organismal populations from the same species or close related species (as suggested in Fig 3 to 7) by means of direct sum of their group representation into Abelian cyclic groups. A general illustration of this modelling would be, for example:

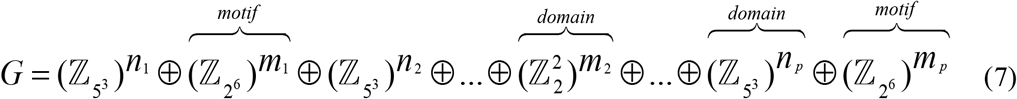

where the exponents *n_i_* and *m_i_* stand for the number of components in the given homocyclic group. That is, Eq. 7 expresses that any large enough genomic region can be represented as direct sum of homocyclic Abelian groups of prime-power order. In other words, the fundamental theorem of Abelian finite groups (FTAG) has an equivalent in genomics.

#### Theorem 1.

The genomic architecture from a genome population can be quantitatively represented as an Abelian group isomorphic to a direct sum of homocyclic Abelian groups of prime-power order.

The proof of this theorem is self-evident across the discussion and examples presented here. Basically, group representations of the genetic code lead to group representations of local genomic domains in terms of cyclic groups of prime-power order, for example, 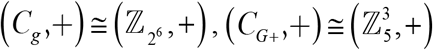 or 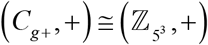, till covering the whole genome. As for any finite Abelian group, the Abelian group representation of genome populations can be expressed in terms of a direct sum of Abelian homocyclic groups of prime-power order. Any new discovering on the annotation of a given genome population will only split an Abelian group, already defined on some genomic domain/region, into the direct sum of Abelian subgroups.

The application of the FTAG in terms of the group representation of genomic regions *G*, as given in Eq. 7, establishes the basis to the study the molecular evolutionary (mutational) processes in terms of endomorphisms. That is, fixed mutational events in the organismal population can be modeled as homomorphism: endomorphisms and automorphisms, all elements of the endomorphism ring 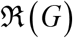 on *G* (see next section). In the context of comparative evolutionary genomics, the analysis of the endomorphism ring 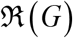 is an intermediate step for the further application of methods from Category theory, which has the potential to unveil unsuspected features of the genome architecture, hard to be inferred from the direct experimentation.

### 3.2 The endomorphism ring

A biologically relevant application of the theory presented here relies on the fact that if a finite group *G* is written as a direct sum of subgroups *G_i_*, as given in Eq. 7, then the endomorphism ring *End*(*G*) is isomorphic to the ring matrices (*A_ij_*), where *A_ij_* ∈ *Homo*(*G_i_*, *G_j_*) (homomorphism between *G_i_* and *G_j_*), with the usual matrix operations [35]. In the case of genomic regions from the same species or closed related genomic regions from distinct species, the endomorphism that transform the DNA aligned sequence *α* into *β* (*α*, *β* ∈ *G*) is represented by a matrix with only non-zero elements in the principal diagonal. These diagonal elements are sub-matrices *A_ii_* ∈ *End*(*G_i_*) or *A_ii_* ∈ *Aut*(*G_i_*). In other words, mutational events fixed in gene/genome populations can be quantitatively described as endomorphisms and automorphisms.

In the Abelian *p*-group defined on 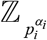, the endomorphisms 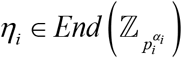 are described as functions *f*(*x*) = *k x* mod 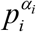, where *k* and *x* are elements from the set of integers modulo 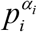. For example, in the cube ACGT the sequence ATACCCATGGCCAAC (blue block in Fig. 3) represented by the vector 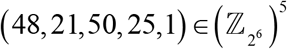 is transformed into the sequence ACACCCATGACCAAC, represented by the vector 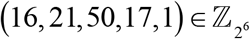, by the automorphism:

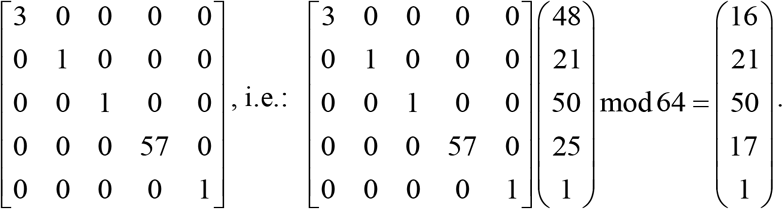

Now, it is not difficult to realize that the set of all endomorphisms 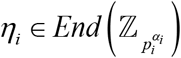 hold the ring axioms given in Appendix A. That is, the set of all endomorphisms 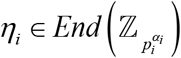 forms a ring on 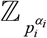 that we shall denote as 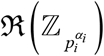.

As shown in reference [35], if *G* = *G*_1_ ⊕ *G*_2_… ⊕ *G_n_ is a direct decomposition with fully invariant summands*, then:

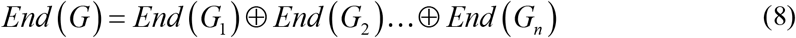

In this modeling, mutational events are represented as endomorphisms 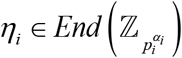 on 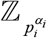. This fact permits the study of the genome architecture through the study of the evolutionary (mutational) process in a genome population. Moreover, the decomposition of the endomorphism ring into subgroups, quotient groups, and cosets can lead to a deterministic algebraic taxonomy of the species based on their genome architecture, which is not limited by our current biological knowledge.

Particularly relevant for the evolutionary comparative genomics is Baer-Kaplansky theorem: *If G and H are p-groups such that* 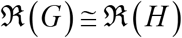, then 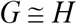 ([34,47]). That is, two Abelian finite groups are isomorphic if, and only if, their endomorphism rings are isomorphic [47]. In other words, genomic regions experiencing mutational events representable by isomorphic rings are algebraically represented by isomorphic Abelian groups and, consequently, have similar genome architecture.

Application of Baer-Kaplansky theorem implies that two gene-body regions encoding exactly for the same polypeptide but with different region architecture (Fig 5) are under different evolutionary pressure. That is, if the group representations of two homologous gene-body regions are not isomorphic, then their endomorphism rings are not isomorphic either and, consequently, they will be under different evolutionary pressure, experiencing different subsets of mutational events, which are represented as endomorphisms from their corresponding endomorphism rings. This scenario is typically found in some isoforms, which are proteins that are similar to each other and perform similar roles within cells [48]. This is the case where two or more closely related genes are responsible for the same translated protein, illustrated in Fig 5. They can be simply duplicated, or paralogous genes, where both paralogs can remain similar (paralog isoforms) if an increased production of the protein is advantageous or if a dosage balance occurs in conjunction with other gene products or where different transcripts can lead to different subcellular localization [49].

A screening of mutational events on subsets of aligned genes suggests that the decomposition of protein-coding regions is tractable, conforming Eq. 8. Results with the alignments of several protein-coding regions are shown in Fig 8. In this example, we searched for automorphisms on the *groups of dual cubes* [11]: ACGT – TGCA and CATG – GTAC on 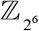, which comprise four of the 24 possible algebraic representations of the standard genetic code [20] isomorphic to 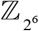.

The analysis of the frequency of mutational events (automorphisms, COVID: human *vs* bat strains) by mutation types is shown in Fig 8**a**. Results are consistent with the well-known observation highlighted by Crick: *the highest mutational rate is found in the third base of the codon, followed by the first base, and the lowest rate is found in the second one* [50]. However, estimations on different gene sets suggest that the evolutionary pressure on each codon position depends on the physicochemical properties (annotated according to IUPAC nomenclature [36]) of DNA bases. For example, in Fig 8**a** pyrimidine (Y) transitions on the third codon position (HHY) are, by far, the most frequent observed mutational events. While, in BRCA1 gene (SI Fig 2), the frequency of purine (HHR) transitions is followed by pyrimidine (HHY) transitions.

The analysis on the pairwise alignment of protein-coding regions of SARS and Bat SARS-like coronaviruses is presented in Fig 8**b** an **c**. Most of the mutational events distinguishing human SARS from Bat SARS-like coronaviruses can be described by automorphism on cube ACGT. This observation was confirmed in primate somatic cytochrome c (Fig 8**c**) and BRCA1 DNA repair gene (Fig 8**d**). Since automorphisms transform the null element (gap-triplet DDD/---) into itself, insertion-deletion mutational events cannot be described by automorphisms but as translations on the groups (denoted as *Trnl* in Fig 8). The representation of conserved genomic regions with homocyclic *p*-group is straightforward. However, their frequency in the genome architecture exponentially decreases with the size of the region (Fig 8**f** and SI Fig 4).

Next, under the assumption that Eq. 8 holds, different protein-coding regions must experience “*preference*” for specific type of automorphisms. To illustrating the concept, an analysis based on the application of Theorem 1 and Eq. 8 on gene/genome population studies, an application of decision tree algorithms was conducted on primate BRCA1 genes. Results for the analysis with Chi-squared Automated Interaction Detection (CHAID) is presented in Fig 9. It is important to keep in mind that this is only an illustrative example with small sample size, and that definite conclusions related to BRCA1 genes can only be derived with larger sample size from humans and non-human primate sequences. In this algorithmic approach, for each compound category consisting of three or more of the original categories, the algorithm finds the most statistically significant binary split for a node (split-variable) based on a chi-squared test [52].

For a given MSA of protein-coding regions, the resulting decision tree leads to stochastic-deterministic logical rules (propositions) permitting a probabilistic estimation of the best model approach holding Eq. 8. For example, since only one mutational event human-to-human from class A3 is reported in the right side of the tree (Fig 9), with high probability the proposition: “(A4 ˅ (A3 ˄ ¬ HRH) → ¬ human” is true. That is, with high probability only non-humans hold the last rule. Due to graphic printing limitations not all tree details are shown in Fig 9 (calculations details are given in the tutorials links provided at the Supporting Information (SI)).

Results shown in Fig 9 were limited to small sample data set and to the application of a relatively “modest” (unsupervised) machine-learning approach which, however, is sufficient to illustrate the concepts. Next, let us suppose that the decision tree from Fig 9 holds on a large enough sample-size (to minimize the classification error) of primate BRCA1-gene populations. Then, with high probability the logical rule: “A1 ˄ R3 ˄ (YHH ˅ HHY) → human” is true. That is, with high probability transitions mutations (T ↔ C) on region R3 from BRCA1 gene (specifically at positions 323 and 333, Fig 9) in the first and third codon positions, represented by automorphisms with coefficient between 0 and 15, are not observed in primates other than humans.

Obviously, the predictive power of the stochastic rules depends on the size of the samples from the populations under scrutiny. A larger data set including 41 variants of the BRCA1 gene and a rough estimation of the (encoded) *mutational cost* given in the term of a quasichemical energy of aminoacid interactions in an average buried environment [11,53] (data included in the *GenomAutomorphism* R package [33]) allow reaching more robust rules after the application of decision tree algorithms. Likewise, an estimation of *mutational cost* can be given in terms of distances between aminoacids based on codon distances defined on a specific genetic-code cube model or on a combination of two models [11,54]. Examples of stochastic some mutational rules are given in Table 1.

**Table 1.** Examples of stochastic mutational rules found in aligned DNA sequences from primate BRCA1 genes.

| Mutational cost (MC) | Stochastic Rule<sup>3</sup> |
| --- | --- |
| Aminoacid contact potential<sup>1</sup> | MC(0.03) $\rightarrow \neg$ human |
| | MC(-0.47) $\wedge$ R4 $\wedge$ A4 → human |
| | MC(-0.47) $\wedge$ (R0 $\vee$ R0. $\vee$ R3 $\vee$ R5) → bonobos |
| | MC(0.08) → bolivian monkey |
| Aminoacid distance based on genetic-code codon distances<sup>2</sup> | MC(1.34) $\wedge$ R0 $\rightarrow \neg$ human |
| | MC(1.36) → gorilla |
| | MC(0.28) $\wedge$ R4 $\wedge$ A4 → human |
| | (MC(0.12) $\vee$ MC(0.12)) $\wedge$ (R1 $\vee$ R5) → silvery gibbon |
| | MC(0.26) $\wedge$ (A1 $\vee$ A2) $\wedge \neg$ R4 $\wedge \neg$ HHW → human |
| | MC(0.99) $\wedge$ HHS $\wedge$ (R7 $\vee$ R4) → golden monkey |

<sup>1</sup>Aminoacid contact potentials are given in reference [53]. <sup>2</sup>Aminoacid distance based on the codon distance are given in reference [54] and applied (together with the concept of encoded mutational cost) in reference [11]. <sup>3</sup>The decision trees using CHAID algorithm are given in the Supporting Information (also available in the tutorials at https://genomaths.github.io/genomautomorphism).

Our results provides supporting evidence to the previous finding reported in [11] about that the selection of the genetic-code cube model cannot be arbitrary, since the automorphisms and the estimation of mutational costs (as defined in [11]) on different local DNA protein-coding regions shows clear “preference” for specific models. Obviously, the mathematical model is only a tool (a representation of the physicochemical relationships given between molecules) applied to uncovering the existence of specific evolutionary constraints for the transmission of genetic information.

### 3.3 Future theoretical developments

In this section we want to highlight a direction of future theoretical developments. A full coverage of this topic is out of the limits of the current work. Nevertheless, a sketch on a future direction is presented here. Our goal will be the description of mutational process on protein-coding regions in terms of homomorphisms of different algebraic structures.

Genomic regions represented as an Abelian group decomposable into homocyclic Abelian *p*-groups, e.g. 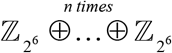, can be studied as *R*-algebras [21], which in particular is a *R*-module and after considering only the sum operation of the ring 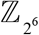, it is also a *G*-module. Recall that our modeling just takes advantage of the group isomorphism: 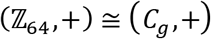 (for the sake of simplicity we are using the same sum operation symbol in both groups, 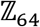 and *C_g_*). Thus, the 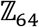-algebra of the group 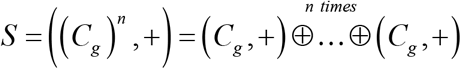 over the ring 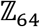 can be defined [21].

In our current case (considering the codon coordinate level), we are interested on heterocycle groups *S* = ⊕*G_i_* of *C_g_* and *C*_*g*+_ (*G_i_* ∈ {*C_g_*, *C*_*g*+_}), as suggested in Fig. 1, which permits the analysis of multiple sequence alignments including insertion-deletion (indel) mutations. It is not hard to notice that the collection of all the *R*-Module of groups S over the ring 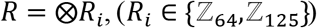 together with *R*-Module homomorphisms conform to a category of ***R*-Modules**, also denoted as ***R*-Mod**. Let 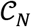 be the category **Ab** with the Abelian groups of the DNA sequences of length ≤*N* as objects and group homomorphisms as morphisms (see Appendix A). Fredy’s theorem states that every Abelian category is a subcategory of some category of modules over a ring [55]. Mitchell has reinforced Fredy’s result, proving that every Abelian category is a full subcategory of a category of modules over a ring [56].

At codon coordinate level, the group defined on the set of codon is a subgroup of the group defined on the set of extended base-triplets (*C_g_* ⊂ *C*_*g*+_) and the 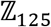-Module of group *C_g_* is a submodule of the 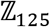-Module of group *C*_*g*+_ over the ring 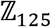. The triplet of gaps ‘---’ corresponds to the identity element of group *C*_*g*+_, which is mapped into 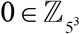 by 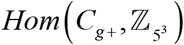. A homomorphism always maps the identity element from the domain of group, say **0**_*C_g_*_, into the identity element from the codomain **0**_*C*_*g*+__, which in *C*_*g*+_ is 0_*C*_*g*+__ = ‘---’.

The following example illustrates a possible sequence of attainable analytical steps with concrete computational biology application. Let *A* = GACAGAGCAGTATTAGCTTCACAC and *B* = GAAAACGTATTATCAAAG DNA sequence segments represented as elements from the groups: 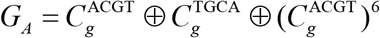 and 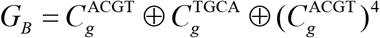, respectively, where 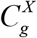 is the Abelian *p*-group defined on the set of 64 codons and base orders (cubes): *X* = {ACGT, TGCA}. Groups *G_A_* and *G_B_* are elements of the **Ab** category 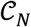 defined on the collection of heterocyclic group 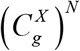 defined on the set of DNA sequences (of codons) with length *N*≤ 8.

Since the triplet of gaps cannot be arbitrary allocated in the sequence, the alignment of DNA sequence is an essential step required for the application of this modeling preserving the biological meaning. The pairwise alignment of the corresponding aminoacid sequences from *A* and *B* yields: 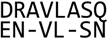, which corresponds to the DNA sequence alignment: 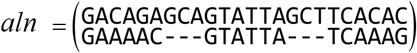. That is, to preserve the reading frame, a robust alignment is accomplished translating the codon sequence into aminoacid sequence alignment.

Sequences *A* and *B*’ = GAAAAC---GTATTA---TCAAAG can also be represented as elements from group:

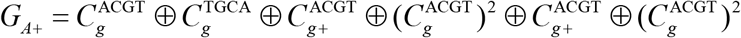

This group is an element of the **Ab** category 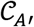, which is a subcategory of the ***R*_*A*+_-Mod** category 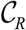 over the ring 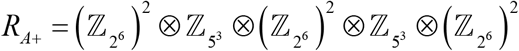. The group homomorphism *F_B_*: *G_B_* → *G_B′_* is a functor that maps DNA sequences from group 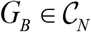 into an element from group 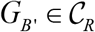 (see Appendix B). That is, for all element *b* = (*X*_1_,*X*_2_,*X*_3_,*X*_4_,*X*_5_,*X*_6_) (*b* ∈ *G_B_*) there is a unique element 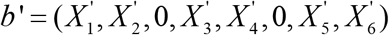 (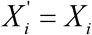 and *b*′ ∈ *G_B′_*, ⊂ *G_A′_*).

Also, there is an injective morphism *F_A_*: *G_A_* → *G_A′_*, that transforms each element *a* = (*X*_1_,*X*_2_,*X*_3_,*X*_4_,*X*_5_,*X*_6_,*X*_7_,*X*_8_) (*a* ∈ *G_A_*) into a unique element 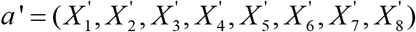 (*a*′ ∈ *G_A′_*), which is evident since *C_g_* ⊂ *C*_*g*+_ and, consequently, codons are preserved, i.e., 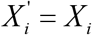, and *G_A_* is isomorphic to the image *F_A_*(*G_A_*). The homomorphism *F*_*A*+_: *G_A_* → *G*_*A*+_ is also a functor which maps elements from the ***R_A_*-Mod** category over the ring 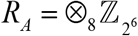 into the ***R*_*A*+_-Mod** category. Notice that *F_B_*(*G_B_*) is a subgroup of *F_A_*(*G_A_*).

In practice, for the sake of computational genomics implementations, the aligned DNA sequences *A* and *B* can be represented by the numerical vectors a *a* = (9,32,24,56,60,27,28,5) and *b* = (8,1,56,60,28,2), respectively, with coordinates on the ring ***R*_*A*+_**. The coordinates of sequences A and B on the ring ***R*_*A*+_** and permits the new representations: 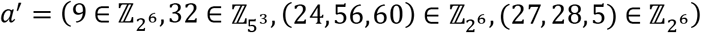 and 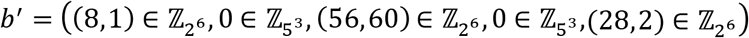, respectively. Sequence *a′* is map into *b′* the group homomorphism *φ*: *a*′ → *b*′:

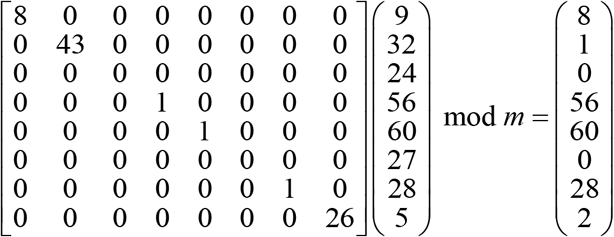

where *m* = (64,125,64,64,64,64,64,64). A group homomorphism *h*: *G_B′_* → *G_A_* to accomplish the mapping *h*(*b*′) = *a*′ is computed as:

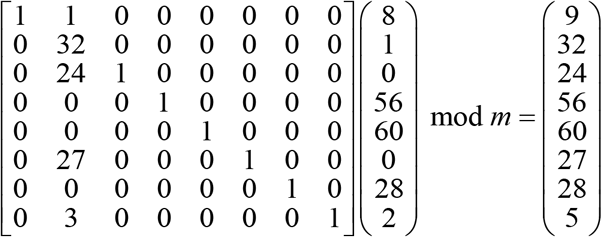

Or by means of the affine transformation:

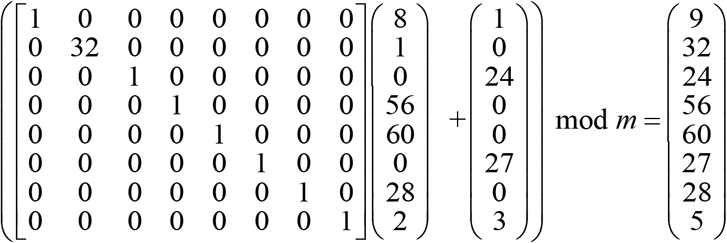

In summary, a future theoretical development in the framework of category theory opens new horizons for the analysis of the mutational process in a wider computational genomic scenario not previously studies in molecular evolutionary biology (see Material and Methods, section 2.3). This computational biology scenario can be translated into concrete Bioinformatics applications [33].

## 4 Discussions

The encoding of the physicochemical relationships between nucleotides (nitrogenous bases) in the DNA double helix in terms of group operations permits a mathematical representation of genome architecture interpretable in a molecular evolutionary context. The group representations of the genetic code are logically extended from protein-coding DNA regions to the entire genome. As shown in Fig 1, the Abelian group representation of genomic regions into the direct sum of Abelian *p*-group is only one of several proposed steps addressed to get better understanding on how genomes are built in an algebraical biology context.

The theory and concretes examples provided here make explicit the basic foundation for a further unprecedented application of the last advances in Abelian group theory incorporating methods from Category theory, where groups and group homomorphisms (in our context: mutational events) are the main players, which have the potential to discover unsuspected features of the genome architecture, opening new horizon to the genomic taxonomy of species in accordance with the state-of-the-art in mathematics, logic, and computational sciences.

For the sake of reader’s comprehension, the examples on the group representation of genomic regions presented here are simple. However, the analysis demands for the development of novel computational algebraic approaches to study the genomic architecture. Unlike to traditional computational algebra, we can take advantage of the group isomorphisms, which permits decreasing the computational complexity by avoiding symbolic computation. Nevertheless, results presented here show that the architecture of genome region in an entire population can be quantitively studied in an algebraical framework.

Examples shown in Figs 3 to 4 indicates that whatever would be the genomic architecture for given species, the observed variations in the individual populations and in populations from closed related species, it can be quantitatively described as the direct sum of Abelian cyclic groups. The discovering/annotation of new genomic features will only lead to the decomposition of previous known Abelian homocyclic or cyclic groups representing a genomic subregion into direct sums of subgroups. In such algebraic representation DNA sequence motifs for which only substitution mutations happened are specifically represented by the Abelian group 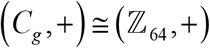, in protein coding regions, and by any combination of groups 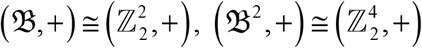 or some quotient group like 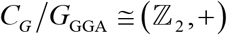 in non-protein coding regions.

Notably, the genetic code Abelian group 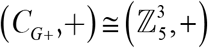 is enough for an algebraic representation of the genome population from the same species or close related species. However, such a decomposition leads to a poor description of local architecture that, as suggested in Figs. 3 to 6, can mask relevant biological features. Figure 3 to 6 illustrate the basic Abelian group representations for further analysis of genome architecture through the study of the mutational events, as essential transformations inherent to the molecular evolutionary process, in terms of endomorphisms and automorphisms, elements of the endomorphism ring.

Particularly promising is the application of the genomic Abelian groups on epigenomic studies, which results from the model where the fifth base *D* stands for the methylated cytosine and adenine. As suggested in Fig 7**a** and **b**, a precise decomposition of methylation motif into the direct sum of Abelian finite groups can lead to their classification into unambiguous equivalence classes. The group structure of the methylation regulatory regions: **GATC**TTTTATGC and GGTTAAAA**GATC**, both represented by the homocyclic group on 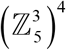, breaks from the monotone homocyclic group representation of the region in terms of cyclic groups on 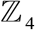 (Fig 7**a** and **b**). The group representation of protein-coding regions (or base-triplet sequences) as numerical vectors with coordinates on 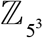

Results indicate that, as a consequence of the genetic code constraints and the evolutionary pressure on protein-coding regions, stochastic-deterministic logical rules can be inferred on a large enough sample-size from a gene/genomic-region population. Such a stochastic-deterministic rules lead to specific applications of Theorem 1 and Eq. 8, consequently, the analysis of mutational process on each group, subgroup, and coset. For example, mutational events on a MSA column (identified) from class YHH (with discriminatory classification power as shown in Fig 8) where the second and third DNA bases remain invariant (H) and the first base are pyrimidines (Y) experiencing transition mutations (across individuals sequences) are represented by automorphisms on a subgroup (from the genetic code Abelian subgroup (*C_G_*, +)) defined on the set {THH,CHH} isomorphic to 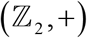 [23].

Figure 9 provides illustrative example that motivates further applications of based machine-learning bioinformatic approaches to unveil the subjacent logic to the genome architecture and its association with the DNA cytosine/adenine methylation patterning found on individual populations and the changes (repatterning) induced by, e.g., environmental changes, aging process and diseases, which is of particular interest in genomic medicine [59]. Machine learning applications on MSA involving large sample size of genomic regions from populations of different species can unveil further decompositions into the direct sum of Abelian groups, which do not depend on our current knowledge of the annotated genomes.

As suggested in Fig 9, we can expect that most of the hiding genomic DNA sequence motif can be unveiled by studying the molecular evolutionary (mutational) process in a genome population through the lens of the endomorphism ring. On this scenario the analysis of group homomorphisms together with decision tree (machine-learning) algorithms permit us the uncovering of stochastic rules constraining the local architecture on genes and genomic regions. In other words, as a consequence of the injective relationship: *DNA sequence* → *3D chromatin architecture* [3,4,6], fixed mutational events (in organismal populations) on DNA sequence motifs involved in the 3D chromatin architecture are under evolutionary pressure, biophysically and biochemically constrained to preserve the chromatin biological functions. The goal is unveiling hidden genomic architecture and rules hard to be detected by current experimental approaches. All the information required can be retrieved from the MSA of DNA sequences, which is particularly relevant for poorly annotated genomes.

Results shown in Figs. 8 and 9 also suggest deep implications of Baer-Kaplansky theorem on the genome architecture. Concretely, on an evolutionary context, the fact that two genomic regions from two different species are identical or almost identical, and that they can even encode the same functional protein, does not necessarily imply that they hold to the same genome architecture. Baer-Kaplansky theorem implies that to hold the same architecture, these hypothetical regions must also experience equivalent evolutionary pressure, which in turn implies that the regions must experience the same type of mutational events in terms of automorphism/endomorphism representations.

For example, let’s suppose that the results shown in Fig 9 were derived from a large sample size (large enough to derive statistically significant rules), then the rule “A1 ˄ R3 ˄ (YHH ˅ HHY) → human” (Fig 9) implies that the gene regions of BRCA1 from human and non-human primates do not belong to the same equivalent class of genomic region. In particular, since the endomorphism rings 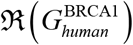 and 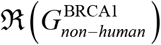 on the Abelian groups 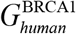 and 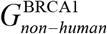 defined on the human and non-human primates BRCA1 genes, respectively, are not isomorphic, then according to the Baer-Kaplansky theorem groups 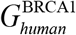 and 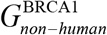 are also not isomorphic. Hence, region architectures of BRCA1 gene in human and non-human primates are (in this hypothetical scenario) implicitly different, which is not obvious to human eyes from their MSA (see supporting information).

Results presented here would have considerable positive impact on current molecular evolutionary biology, which heavily relies on *ad hoc* subjective evolutionary null hypotheses about the past. As suggested in reference [11], the genomic Abelian groups open new horizons for the study of the molecular evolutionary stochastic processes (at genomic scale) with relevant biomedical applications, founded on a stochastic-deterministic ground, which only depends on the physicochemical properties of DNA bases and aminoacids. In this scenario, the only molecular evolutionary hypothesis needed about the past is a fact, the existence of the genetic code.

From the examples provided here, it is clear that there exists a language for the genome architecture unveiled when represented it in terms of sums of finite Abelian groups. Application of Baer-Kaplansky theorem and further studies applying methods of Categorical theory (the potential success of which has been proven in programming languages and in linguistic [57]) can help to unveil the grammar embedded in the DNA sequences. In other words, these applications have the potential to elevate the genomic studies from the current descriptive level to the vanguard level marked in the frontier of science by mathematics, physics, and computational sciences.

Remarkably, further studies applying the theory presented here do not require for special experimental datasets but for the DNA sequences of the genomic regions under scrutiny. Although the accuracy of the predictions depends on the sample size, the number of sequenced genomes stored in the databases grows year-after-year.

## 5 Conclusions

Results to date indicate that the genetic code and the physicochemical properties of DNA bases on which the genetic code algebraic structure are defined, has a deterministic effect, or at least partially rules on the current genome architectures, in such a way that the Abelian group representations of the genetic code are logically extended to the whole genome. In consequence, the fundamental theorem of Abelian finite groups can be applied starting from genomic regions till cover whole chromosomes. This result opens new horizons for further genomics studies with the application of the Abelian group theory, which currently is well developed and well documented [35,60].

Results suggest that the architecture of current population genomes is quite far from randomness and obeys stochastic-deterministic rules. The nexus between the Abelian finite group decomposition into homocycle Abelian *p*-groups and the endomorphism ring paved the ways to unveil unsuspected stochastic-deterministic logical propositions ruling the ensemble of genomic regions and sets the basis for a novel algebraic taxonomy of the species, which is not limited by our current biological knowledge.

In the context of evolutionary comparative genomics, the theory presented here open new horizons for the application of Group theory including methods of Category theory, which have the potential to unveil hidden features and rules inherent to the genome architecture, leading to an unprecedented understanding on how genomes are built.

We believe that the mathematical formalism proposed here sets the theoretical ground for a further development in genomics, transitioning the field from a fully empirical science to a predictive science, where the theoretical and empirical research coexist in a tight positive feedback loop, a development stage only reached so far in the field of physics.

All our claims are feasible, only limited by our computational power and the availability of samples of sequenced genomes from the same species and from multiple species. At this point we emphasize that *an accurate understanding of the genome architecture and population’s structure, on a formal mathematical framework, is as essential for the future of genetic/genome engineering as the physics of architecture is to the design of sturdy and stable energy-efficient buildings*.

## 6 Appendix A. Genetic code algebraic structures defined on the base and codon sets

An Abelian group structure (*B*, +) is a set ***B*** together with a binary operation ‘+’ that combines any two elements *a* ∈ *B* and *b* ∈ *B* to form another element of *c* ∈ *B*, denoted *a* + *b* = *c*, which satisfy the following axioms:

1. Associativity. For all *a*,*b*,*c* ∈ *B*, the equality (*a* + *b*) + *c* = *a* + (*b* + *c*) holds.
2. Identity. There exists an element *e* ∈ *B* named identity element of *B*, such that for any *a* ∈ *B*, the equality *a* + *e* = *a* holds.
3. Commutativity. For all *a*, *b* ∈ *B*, the equality *a* + *b* = *b* + *a*

The Abelian groups considered here are finite cyclic groups (*G*, +) isomorphic to the Abelian group defined on the set of integers modulo *n*, denoted as 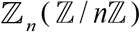. That is, the integers 1,2,3,…,*n* −1 form a cyclic group of order *n* under addition (modulo *n*) and 0 as the identity element. This group will be denoted as 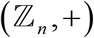. However, for the sake of simplicity in the figures it will be denoted simply as 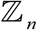, i.e., without making distinction between the set 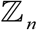 and group structure defined on it. The particular interest for the current work is the Abelian *p*-group derived when *n* = *p^α^* where *p* is a prime number and *a* an integer. The group operations defined on the set of bases or on the codon set are associated to physicochemical properties of DNA bases (see the next sections).

### Homomorphisms and isomorphisms

In modern algebra, a group homomorphism is a map *f*: *A* → *B* between two group structures (*A*,**•**) and (*B*, ∘) such that for all *a*, *b* ∈ *A* holds: *f*(*α*_1_ • *α*_2_) = *f*(*α*_1_) ∘ *f*(*α*_2_) = *β*_1_ ∘ *β*_2_, where *β*, *β*_2_ ∈ *B*. A group isomorphism is a one-to-one correspondence (mapping) between two sets that preserves binary relationships between elements of the sets. That is, an isomorphism is a homomorphism holding the inverse mapping: *f*^−1^(*β*_1_ ∘ *β*_2_) = *f*^−1^(*β*_1_) • *f*^−1^(*β*_2_) = *α*_1_ • *α*_2_. For example, since there exists only one cyclic group with four elements up to isomorphism, for each one of the 24 cyclic group 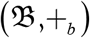 defined on the set of bases 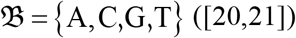 ([20,21]) there exists a one-to-one mapping *f* such that for each base 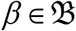 there is an integer 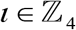 such that *f*(*β*) = *ι* and:

1. *f*(*β*_1_ +_*b*_ *β*_2_) = *f*(*β*_1_) + *f*(*β*_2_) = *ι*_1_ + *ι*_2_, 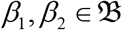 and 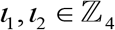.
2. The inverse mapping *f*^−1^(*ι*_1_ + *ι*_2_) = *f*^−1^(*ι*_1_) +_*b*_ *f*^−1^ (*ι*_2_) = *β*_1_ +_*b*_ *β*_2_

To highlight the fact that the sum operations are defined on different ways on the sets 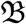 and 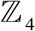, we have used the symbols ‘+_*b*_’ and ‘+’, respectively. However, for sake of brevity of the symbolic notation, such knowledge will be considered implicit, writing simply ‘+’. Then, we said that groups 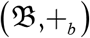 and 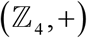 are isomorphic; in symbols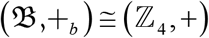. *f* and its inverse *f*^−1^ are called isomorphisms. If *f* (and its inverse *f*^−1^) is a mapping from a group into itself, then *f* is called an automorphism. A mapping *g*, not necessarily one-to-one, of the elements from a group into itself is called a group endomorphism. An endomorphism that is also an isomorphism is an automorphism.

A ring algebraic structure is obtained when together with the sum operation “+” (as defined above) a new operation “·” is defined on the set *B* holding the properties:

1. Associativity: (*a* · *b*)· *c* = *a* ·(*b* · *c*) for all *a*, *b*, *c* ∈ *B*
2. Multiplication is distributive with respect to addition:

a. (*a* + *b*)·*c* = (*a* · *c*) + (*b*·*c*) for all *a*,*b*,*c* ∈ *B* (right distributivity).
b. *c* ·(*a* + *b*) = *c* · *a* + *c* · *b* for all *a*,*b*,*c* ∈ *B* (left distributivity).

As it is shown in the next section, these algebraic structures have been defined on the genetic code. In particular, the ring 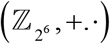 and endomorphism ring (section 3.1) has been defined and studied on the genetic code [21].

## Appendix B. Category

Category theory is a general mathematical theory of structures and of systems of structures that occupy a central position in contemporary mathematics, theoretical computer science, and linguistics [61].

### Definition

A category 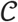 can be described as a collection of objects 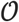 satisfying the following three conditions:

1. *Morphism*: For every pair *X*, *Y* of objects, there is a set *Hom*(*X*, *Y*) called the *morphisms* from *X* to *Y* in 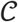. If *f* is a morphism from, we write *f*: *X* → *Y*.
2. *Identity*: For every object *X*, there exists a morphism *id_X_* in *Hom*(*X*, *Y*), called the *identity* on *X* (also denoted as 1_*X*_).
3. *Composition*: For every triple *X*, *Y*, and *Z* of objects, there exists a partial binary operation from *Hom*(*X*, *Y*) × *Hom*(*Y*, *Z*) to *Hom*(*X*,*Z*), called the composition of morphisms in 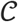. If *f*: *X* → *Y* and *g*: *Y* → *Z*, this composition is written as the mapping (*g* ∘ *f*: *X* → *Z*.

Identity, morphisms, and composition satisfy two axioms:

*Associativity*: If *f*: *X* → *Y*, *g*: *Y* → *Z*, and *h*: *Z* → *W*, then *h* ∘ (*g* ∘ *f*) = (*h* ∘ *g*) ∘ *f*.
*Identity*: If *f*: *X* → *Y*, then *f_X_* ∘ *f* = *f* and *f* ∘; *f_X_* = *f*.

### Definition

A functor *F* is a function between two categories 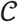 and *D* which maps objects to objects and morphisms to morphisms. That is:

- For each 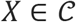 there is an object *F*(*Y*) ∈ *D*
- For each morphism *f*: *X* → *Y* in 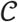 there is morphism *F*(*f*): *F*(*X*) → *F*(*Y*) in *D* such that the following conditions hold:

i. *F*(*g* ∘ *f*) = *F*(*g*) ∘ *F*(*f*) for all morphisms *f*: *X* → *Y* and *g*: *X* → *Y* in 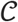
ii. *F*(*id_X_*) = *id*_*F*(*X*)_ for all 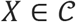.

## 7 Supporting Information

A summary with the reported genetic code Abelian groups relevant for the current study is provided as supporting information in a file named: *Supporting_Information.docx*.

All the data, computational and statistical analyses can be reproduced following the R scripts provided in tutorials available at the GenomAutomorphism R package website https://genomaths.github.io/genomautomorphism/. In particular, data and R scripts used in the computation of automorphisms and the decision tree from Fig 9 are available within GenomAutomorphism R package and in a tutorial at: https://genomaths.github.io/genomautomorphism/articles/automorphism_and_decision_tree.html.

## Supporting information

### 1 Reported genetic code abelian groups relevant for the current study

Herein, we assume that readers are familiar with the definition of abelian group, which otherwise can be found in textbooks and elsewhere including Wikipedia. Nevertheless, all the abelian groups discussed here are isomorphic to the well-known abelian groups of integer module *n*, which are easily apprehended by a college-average educated mind. For example, the abelian group defined on the set {0, 1, 2, 3, 4}, which corresponds to the group of integers modulo 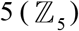, where (2 + 1) mod 5 = 3, (1 + 3) mod 5 = 4, (2 + 3) mod 5 = 0, etc. The subjacent biophysical and biochemical reasonings to define the algebraic operations on the set of DNA bases and on the codon set were given in references [12,14,17].

#### The 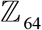 – algebras of the genetic code (C_g_)

The 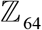 – algebras of the genetic code (*C_g_*) and gene sequences were stated several years ago. In the 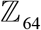 – algebra *C_g_* the sum operation, defined on the codon set, is a manner to consecutively obtain all codons from the codon AAC (UUG) in such a way that the genetic code will represent a non-dimensional code scale of amino acids interaction energy in proteins.

A description of the genetic code abelian finite group (*C_g_*, +) can be found in [12]. Group (*C_g_*, +) is isomorphic to the group on the set 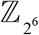 (the sum of integer modulo 64), which formally will be expressed as 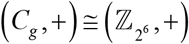. This group on the set 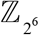 (the sum of integer modulo 64). The mapping of the set of codons *X*_1_*X*_2_*X*_3_ ∈ *C_g_* into the set 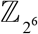 is straightforward after consider the bijection A ↔ 0,C ↔ 1,G ↔ 2,U ↔ 3 and the function *g*(*x*) = 4*x*_1_ + 16*x*_2_ + *x*_3_.

For example:

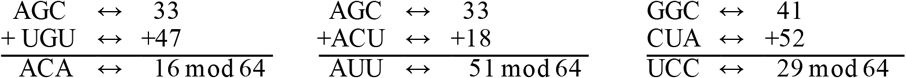

The *Z*_64_-algebra *C_g_*, however, is limited to protein-coding regions, while it is well known that, in eukaryotes, only a small fraction of the genome-about 3%-called open reading frame (ORF) encodes for proteins [18]. Since non-coding DNA sequences can have a base pairs number not multiple of three, complete chromosomes and genomes cannot be described by means of group (*C_g_*,+). In addition, natural genomic variations that includes insertions and deletion mutations (indel mutations) across individuals from the same population and close-related populations from different species cannot be represented with group (*C_g_*,+).

#### 1.1 The 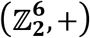 group of the genetic code (C_g_)

Group (*C_g_*, +) is the additive group of a module over a ring, which however, do not conform to a vector space. To build a genetic code vector space, a Galois field (*GF(4*) structure in the ordered base set *B* = {G, U,A,C} was introduced in reference [14]. In particular, an isomorphism with the Galois field is defined by means of its binary representation 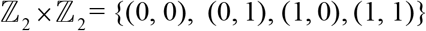, i.e. a unique *GF*(4) up to isomorphism exists, such that a bijection 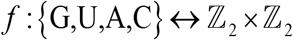 from the DNA base set *B* = {G, U, A, C} to the set of binary duplets (α_1_, α_2_) is stated., where 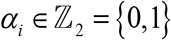. For example, the bijection *f* is defined as:

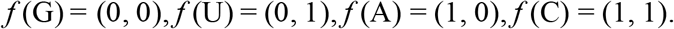

The additive group of bases is the Klein four-group, which is defined by the group presentation: 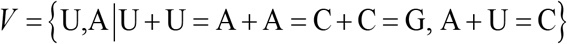, i.e., 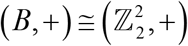. Next, the abelian group on the set of codons *B*^3^ was defined as the direct third power *B*^3^ = *B* × *B* × *B* of the group (*B*, +), i.e. (*B*^3^, +) = (*B*, +) × (*B*, +) × (*B*, +), which is isomorphic to the group: 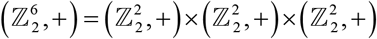, i.e., 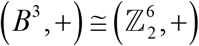. The sum operation on the set (*B*^3^, +) follows from the sum operation by coordinates.

As pointed out before by Crick, the first two bases of codons determine the physicochemical properties of aminoacids [29]. The four encoded amino acids of every class are either the same or show very similar physicochemical properties. This genetic code regularity is captured by the quotient group *B*^3^/*G_GGA_*, where *G_GGA_* is a subgroup of *B*^3^ integrated by the elements {GGG,GGA} (the elements of the quotient group *B*^3^/*G_GGA_* are given in Table 5 from [14]). The quotient group *B*^3^/*G_GGA_* is isomorphic to group 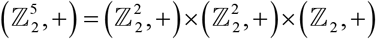. Each element of this group represents an equivalence class of codons. Two triplets *X*_1_*X*_2_*X*_3_ and *Y*_1_*Y*_2_*Y*_3_ are equivalent if, and only if, the difference *X*_1_*X*_2_*X*_3_ + *Y*_1_*Y*_2_*Y*_3_ ∈ *G_DDA_*. In biological terms, substitution mutations involving codons from the same class will not alter (or at least no substantially alter in most of the cases) the physicochemical properties of the encoded protein domains, since in the worst scenario involves aminoacids with very close physicochemical properties, with the exception of codon for aminoacid tryptophan.

#### 1.2 The 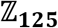 group of the extended genetic code (C_e_)

The extension of the *genetic code group* (*C_g_*, +) follows straightforward from the extension of the codon set, which is easily accomplished extending the source alphabet of the standard genetic code: {A, C, G, U} and, consequently, extending the base triplet set (extended triplet) as *X*_1_*X*_2_*X*_3_, *X_i_*Î{D, A, C, G, U} [22]. The new algebraic structure (*C_e_*, +) is isomorphic to the abelian group defined on the set 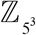 (the sum of integer modulo 125), formally, 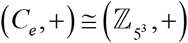. The mapping of the set of codons *X*_1_*X*_2_*X*_3_ ∈ *C_e_* into the set 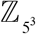 is straightforward after consider the bijection *D* ↔ 0,A ↔ 1,C ↔ 2,G ↔ 3,U ↔ 4 and the function *g*(*x*) = 5*x*_1_ + 25*x*_2_ + *x*_3_ (see Table 1). For example:

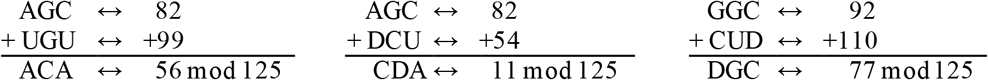

**Table 1.**
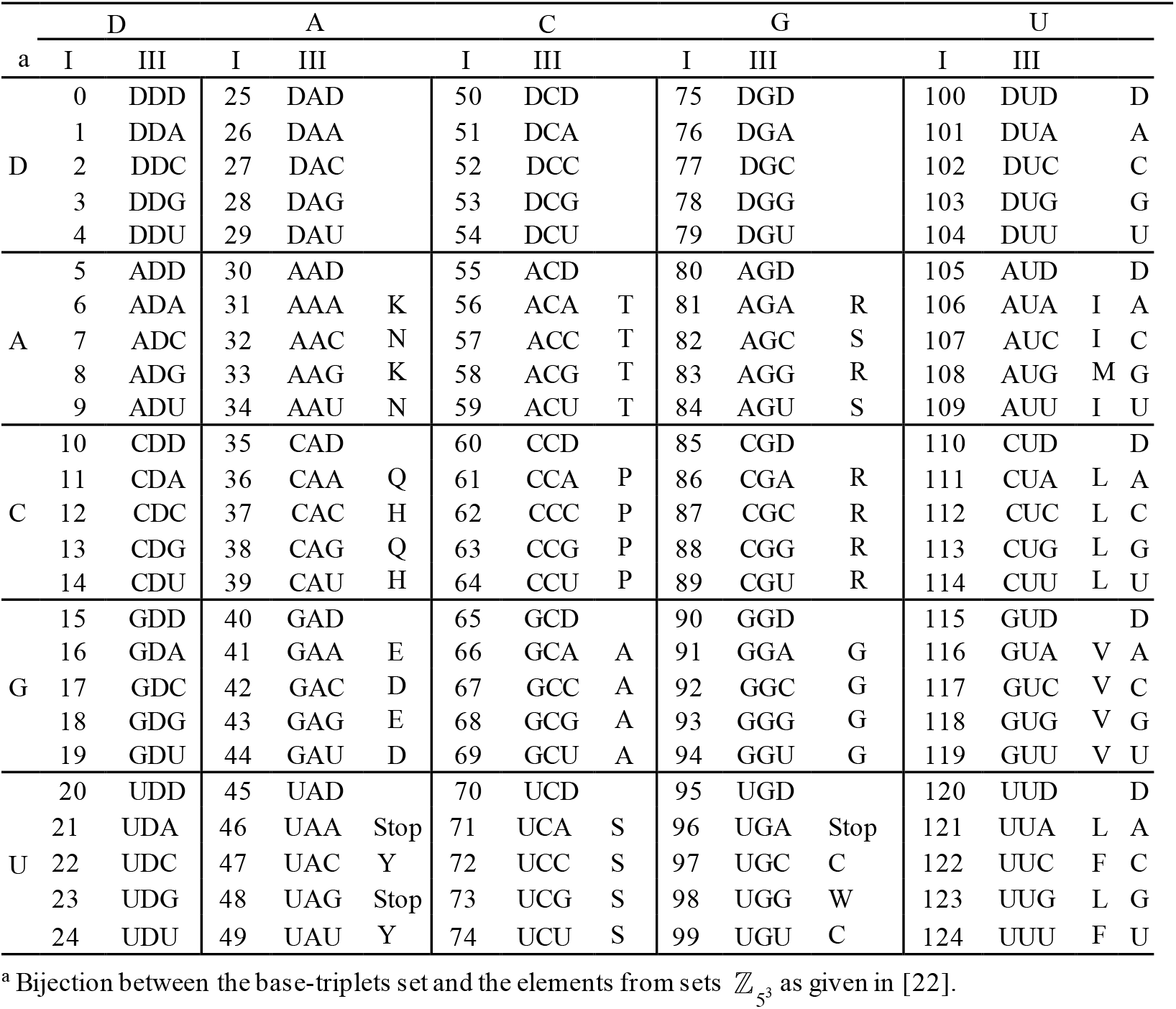
Ordered set of extended triplets corresponding to the elements from 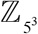

#### 1.3 The 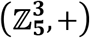 group of the extended genetic code (C_e_)

Formally the genetic code only is limited to translated coding regions where the number of RNA bases is a multiple of 3. However, as suggested in reference [17], the difficulties in prebiotic synthesis of the nucleosides components of RNA (nucleo-base + sugar) and suggested that some of the original bases may not have been the present purines or pyrimidines [18]. Piccirilli et al. [19] demonstrated that the alphabet can in principle be larger. Switzer et al. [20] have shown an enzymatic incorporation of new functionalized bases into RNA and DNA. This expanded the genetic alphabet from 4 to 5 or more letters, which permits new base pairs, and provides RNA molecules with the potential to greatly increase their catalytic power.

It is important to notice that even in the current (*friendly*) environmental conditions not a single cell can survive without a DNA repair enzymatic machinery and that such an enzymatic machinery did not existed at all in the primaeval forms of live. Here, we are confronting the *chicken and egg* problem. To date, the best solution (to our knowledge) is the admission of alternative base-pairs in the primordial DNA alphabet which, as suggested in the studies on the prebiotic chemistry, could contribute to the thermal and general physicochemical stability of the primordial DNA molecules.

Several algebraic structures have been proposed including an additional letter into the DNA alphabet: A, C, G, T. The new letter (D) stands for current insertion deletion/mutations or for alternative wobble base pairing, which would be a relict fingerprint from primordial enzymes derived from a more degenerated ancestral genetic code [17,21,22]. Supporting evidence for the existence of a more degenerated ancestral genetic code built up on a larger alphabet is found in the tRNA anticodon region permitting wobble base pairing by including, e.g., bases such as: inosine (in eukariotes), agmatidine (in archaea), and lysidine (in bacteria), which has been proposed as evolutionary solutions to the need for lower the high translational noise connected to the reading of the AUA and AUG codons [23,24]. Additionally, various alternative base pairs like methylated cytosine and adenine are still present in the current genomes playing an important role in the epigenetic adaptation of organismal populations to the continuous environmental changes [10,25].

Cytosine DNA methylation results from the addition of methyl groups to cytosine C5 residues, and the configuration of methylation within a genome provides trans-generational epigenetic information. These epigenetic modifications can influence the transcriptional activity of the corresponding genes, or maintain genome integrity by repressing transposable elements and affecting long-term gene silencing mechanisms [26,27].

The Galois field *GF*(5) of the DNA set of bases 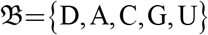 was introduced in reference [17]. This structure led to the definition of a 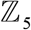 – vector space 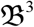 over the set 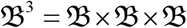 isomorphic to the set 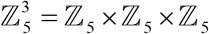 [17,30]. But here, we are interested only in the abelian groups 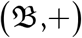 and 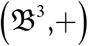. After the bijection *D* ↔ 0, A ↔ 1,C ↔ 2,G ↔ 3,U ↔ 4, the sum operation of two DNA bases follows from the sum operation on the Galois field *GF*(5) (i. e., on 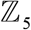, the sum of integers modulo 5). For example, C + U ↔ (2 + 4) mod 5 = 1 ↔ A. The sum operation on the set 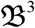 follows from the sum operation by coordinates.

It is worthy to notice that there 24 way to define each one of the above mentioned algebraic structures [30,31]. Nevertheless, for each defined genetic code group, there is only one (genetic code abelian group) up to isomorphism, which lead to their representation as an abelian group, where the sum operation corresponds to the sum of integer modulo *n* ∈{2,2^6^,5,5^3^}.

### 2 REST DNA sequence motifs. R code

The DNA sequence motifs targeted by transcription factors usually integrate genomic building block across several species

DNA sequence alignment of the protein-coding sequences from phospholipase B domain containing-2 (PLBD2) carrying the footprint sequence motif recognized (targeted) by the Silencing Transcription factor (REST), also known as Neuron-Restrictive Silencer Factor (NRSF) REST (NRSF).

#### 2.1 Download methylation data from Github

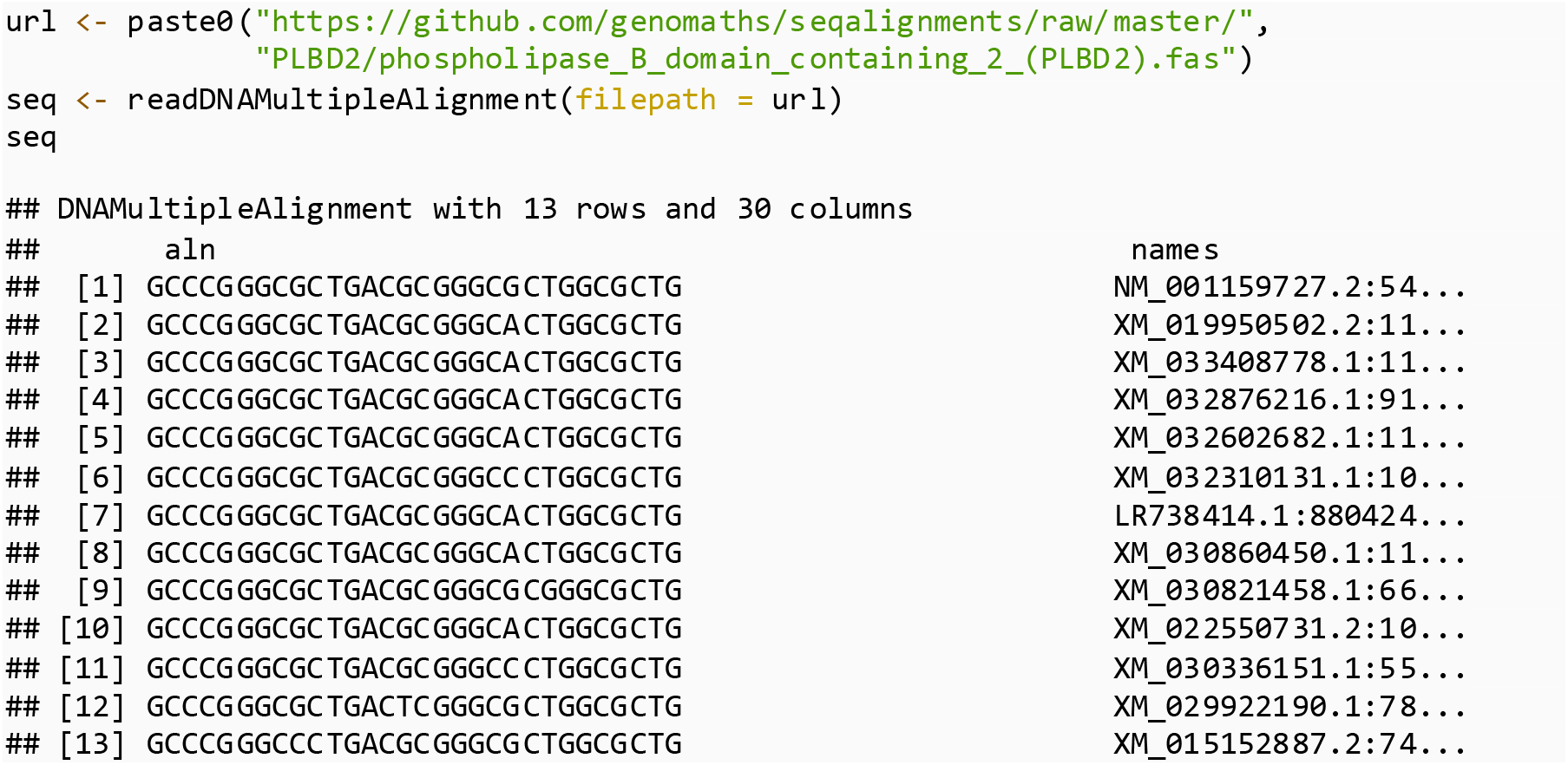

#### 2.2 The Logo sequence

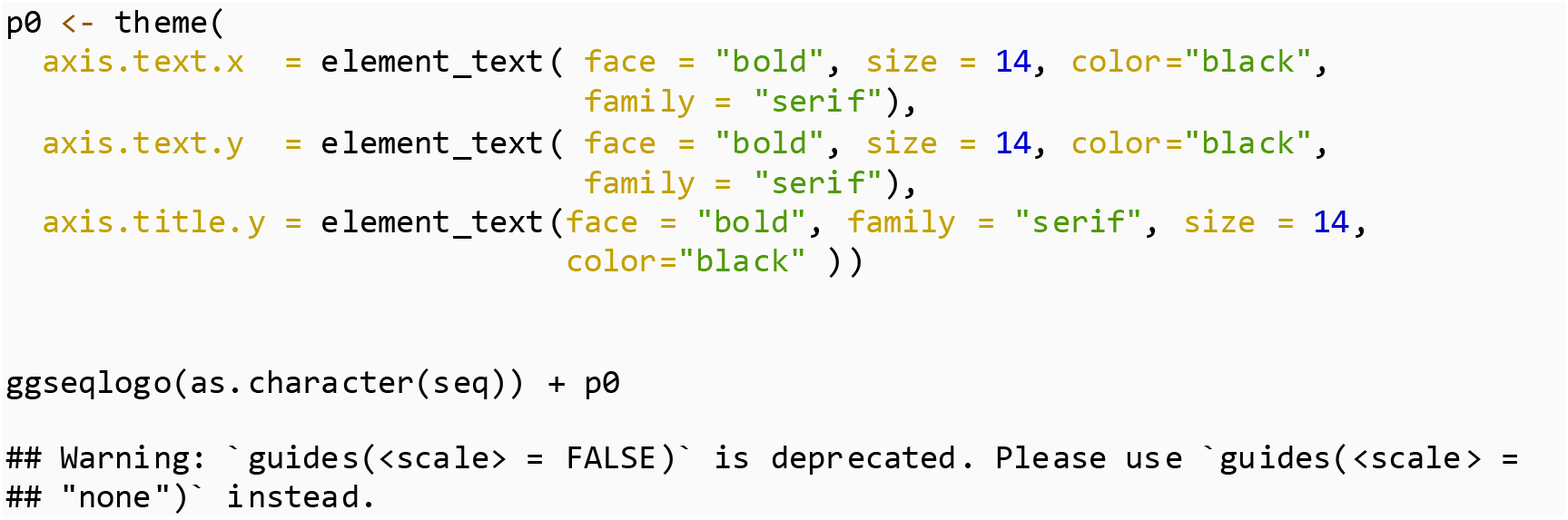

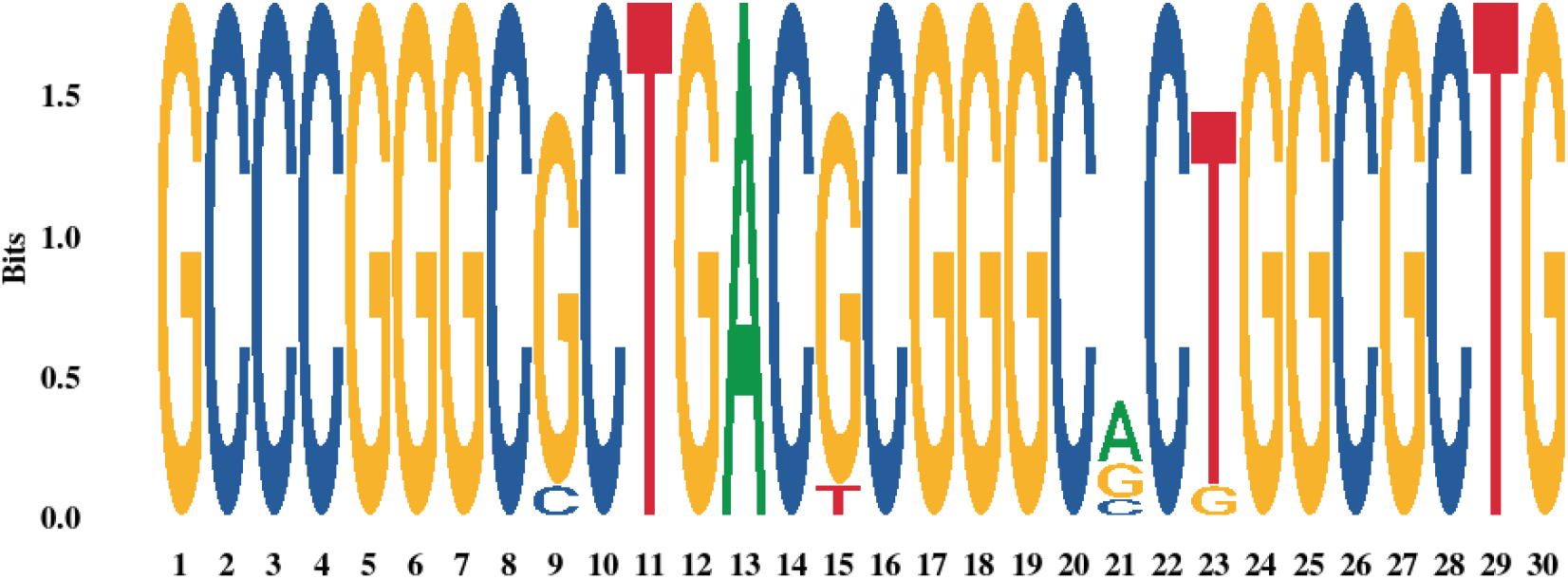

**3 Table S2.** Operation tables of the Klein four group defined on the ordered set of four DNA bases {A,G,C,U}.

| + | A | G | C | U |
|---|---|---|---|---|
| A | A | G | C | U |
| G | G | A | U | C |
| C | C | U | A | G |
| U | U | C | G | A |

**4 Figure. S1.**
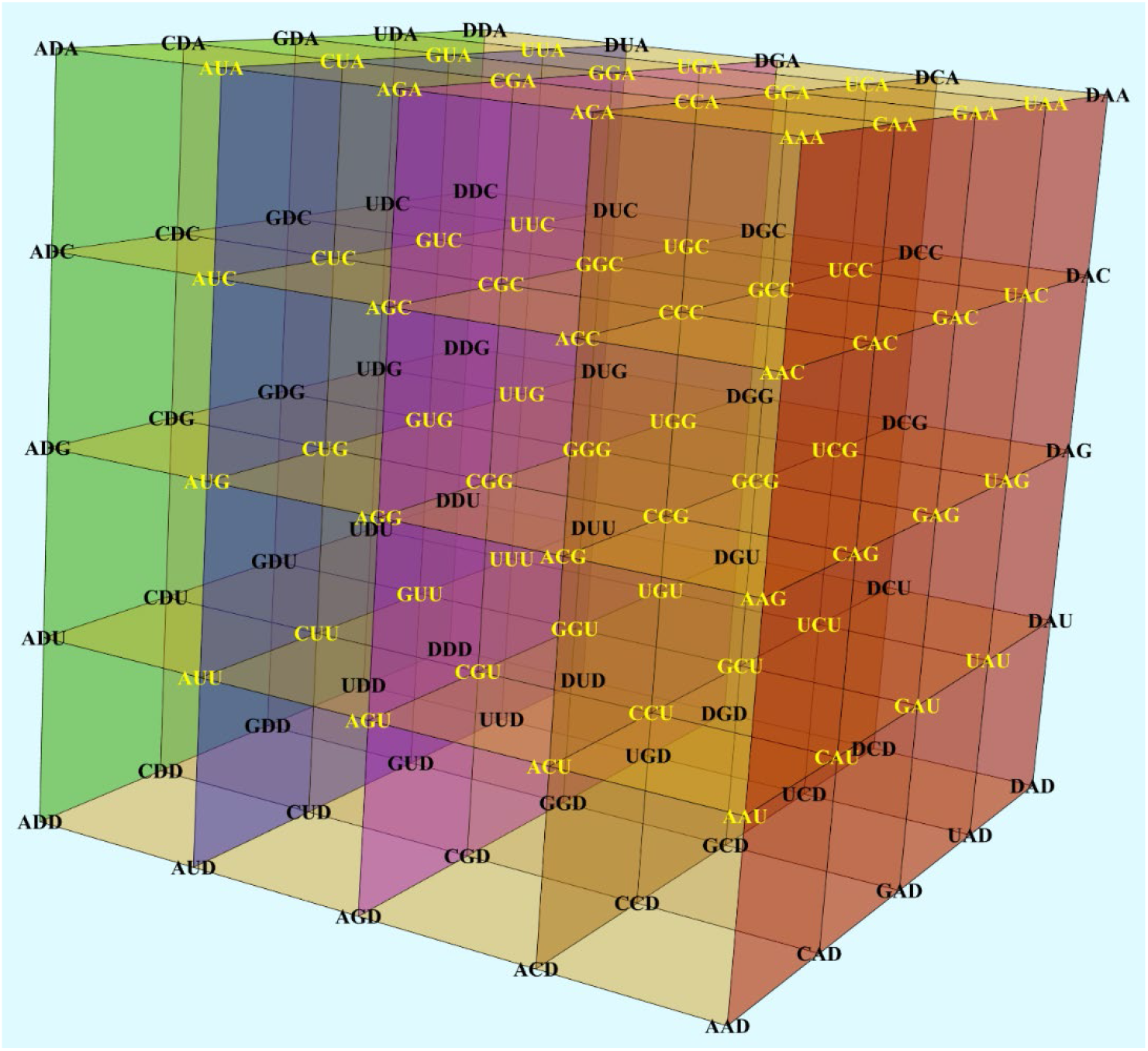
Genetic-code cub UGCA inserted in the three-dimensional space. The insertion of the cube in 3D-space takes advantage of the group isomorphism: 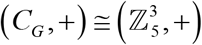, where 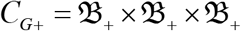 and 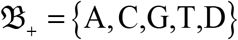. The extended base-triplets including the alternative base D (omitted) are located on the cartesian coordinate planes. Codons (in yellow) encoding for amino acids with similar physicochemical properties are located on the same vertical plane. Cube pair ACGU – UGCA for a group of dual cubes [11].

**5 Figure S2.**
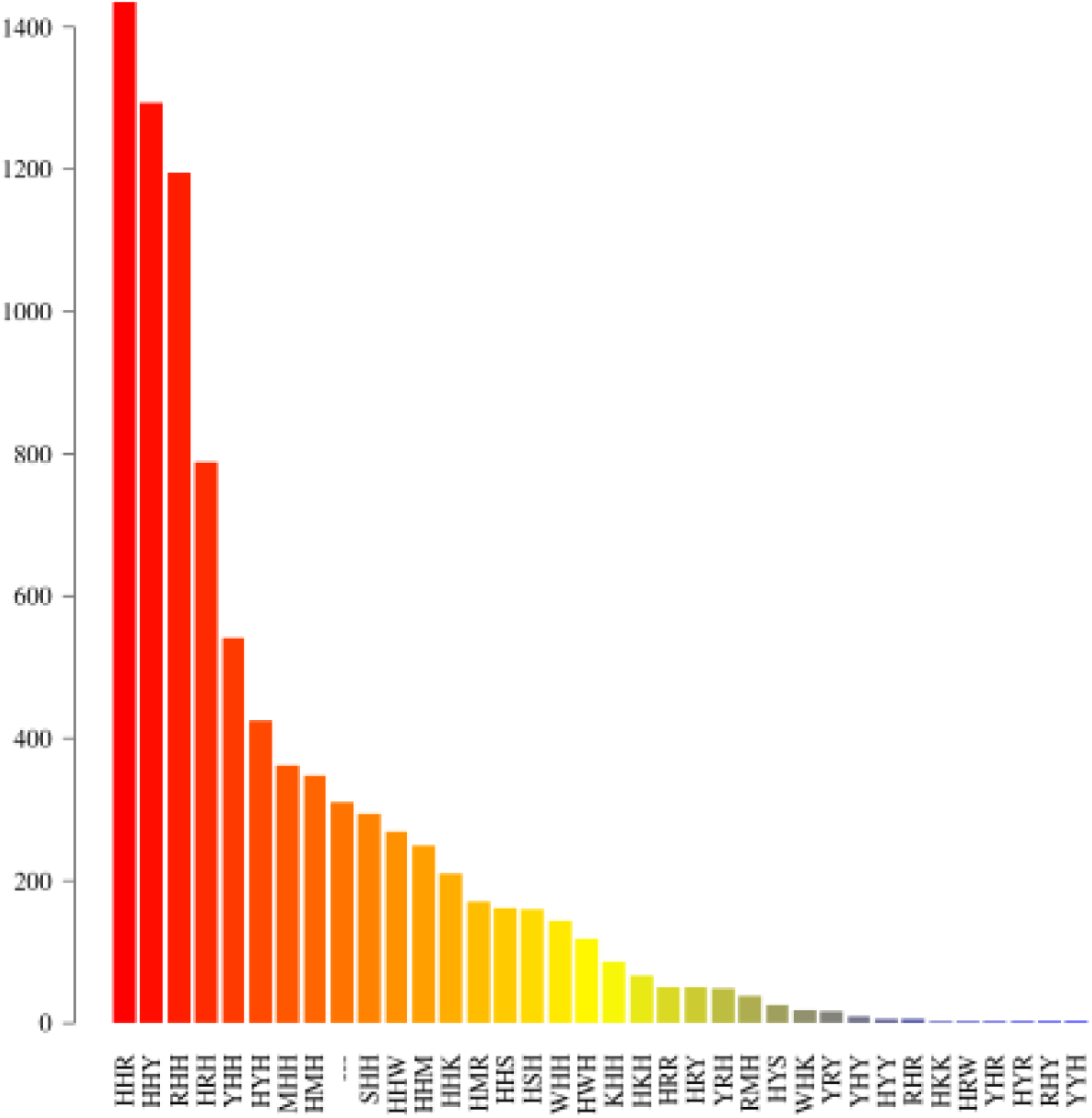
Automorphism distribution per mutation type in primate BRCA1 gene. As a result of the evolutionary pressure on DNA protein-coding regions (addressed to preserve the aminoacid physicochemical properties and, consequently, the biological functions of the encoded proteins) the highest mutational rate is found in the third base of the codon, followed by the first base, and the lowest rate is found in the second one [23]. DNA bases are classified based on the physicochemical criteria used to ordering the set of codons: number of hydrogen bonds (strong-weak, S-W), chemical type (purine-pyrimidine, Y-R), and chemical groups (amino versus keto, M-K) [20]. Preserved codon positions are labeled with letter “H” and insertion-mutations identified in the multiple sequence alignment are labeled as “---”. The data and R script to build this graphic is available at: https://genomaths.github.io/genomautomorphism/articles/automorphism_on_msa_brca1.html.

**6 Figure S3.**
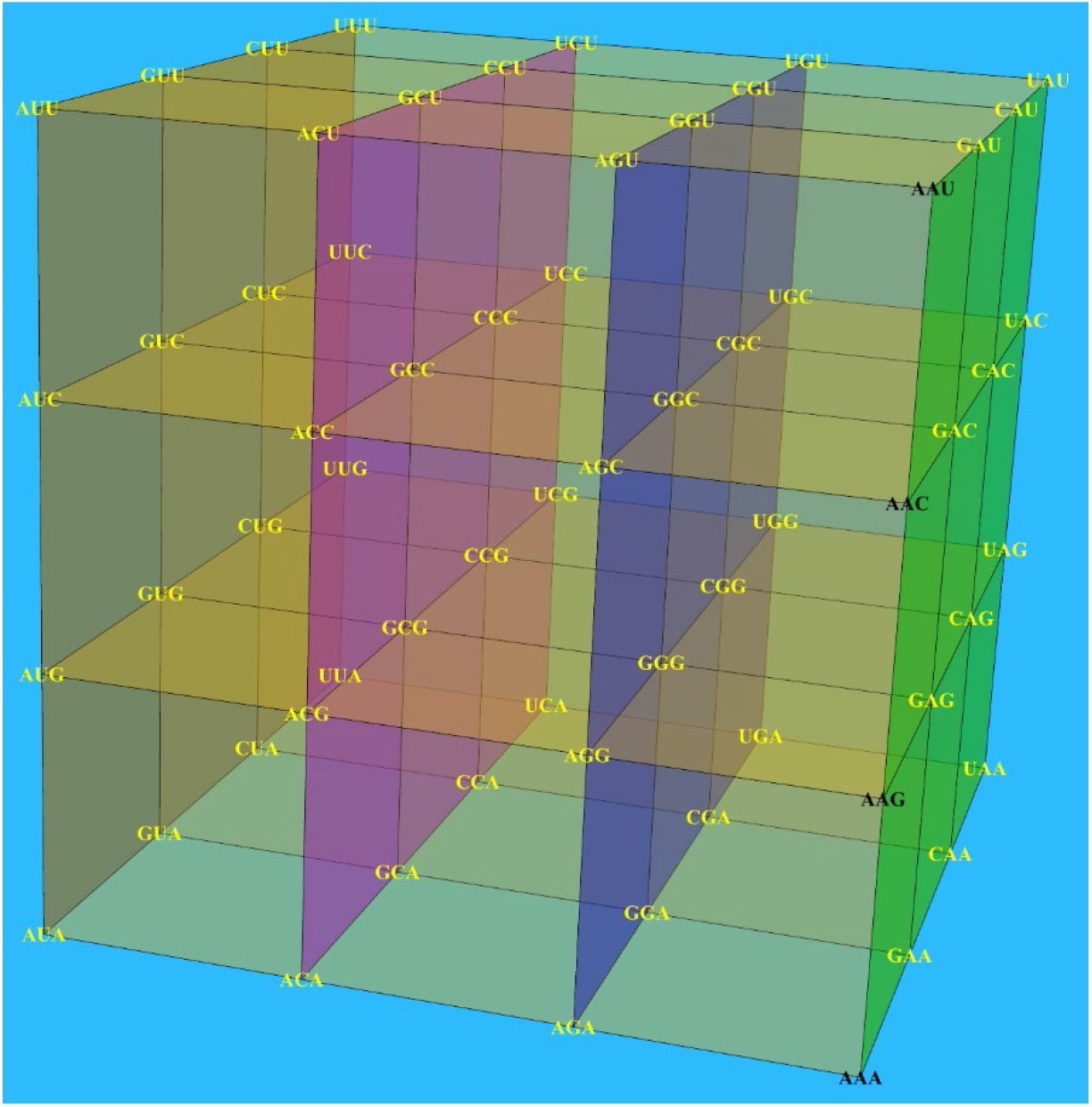
Genetic-code cube AGCU inserted in the three-dimensional space and centered in the coordinate origin. The insertion of the cube in 3D-space takes advantage of the isomorphism: 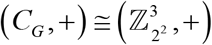, where 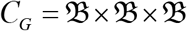 and 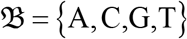. Codons (in yellow) encoding for amino acids with similar physicochemical properties are located on the same vertical plane. The vertical line with codons: AAA, AAG, AAC, and AAU (in black) forms a subgroup isomorphic to Klein four group 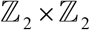. The rest of vertical lines are cosets from the quotient group: (*C_G_*, +)/*G*_AAG_, where *G*_AAG_ = ({AAA,AAG}, +).

**7 Figure S4.**
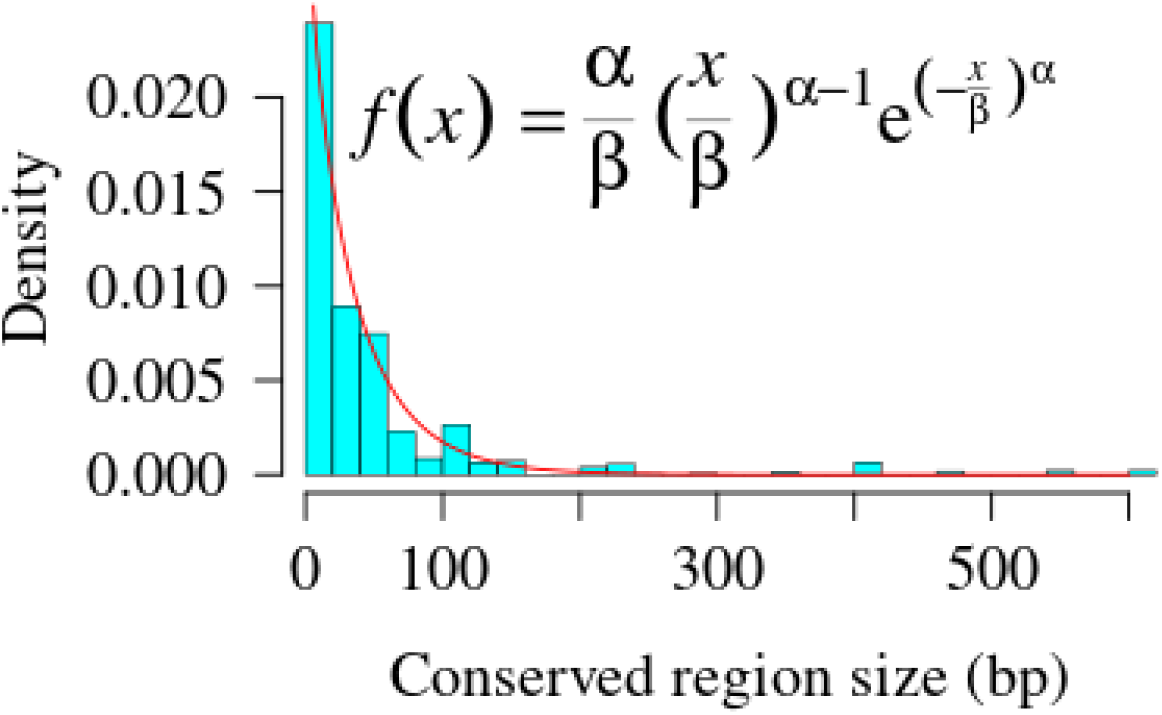
Distribution of the conserved BRCA1 gene from primate according to their size. The best fitted probability distribution turned out to be the Weibull distribution, a member of generalized gamma distribution family.

### 8 Decision tree with a larger dataset

#### 8.1 Using mutational cost in terms of Aminoacid contact potentials

In this analysis a new variable has been include, which derives from the estimation of the contact potential matrix of amino acids made by Miyazawa and Jernigan, which are considered quasichemical energy of interactions in an average buried environment. The amino acids potentials, as well as others numerical indices representing various physicochemical and biochemical properties of amino acids and pairs of amino acids can be found in AAindex. The entire script is available in the tutorial of *GenomAutomorphism* R package [28]: https://genomaths.github.io/genomautomorphism/articles/automorphism_and_decision_tree.html#decision-tree-with-a-larger-dataset

The quasichemical energy of amino acid interactions was stored in *mut_effect* variable

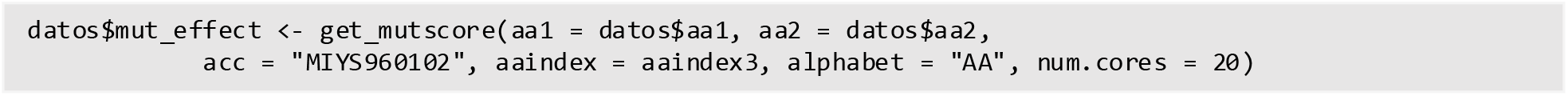

The result from the application of CHID algorithm is given below

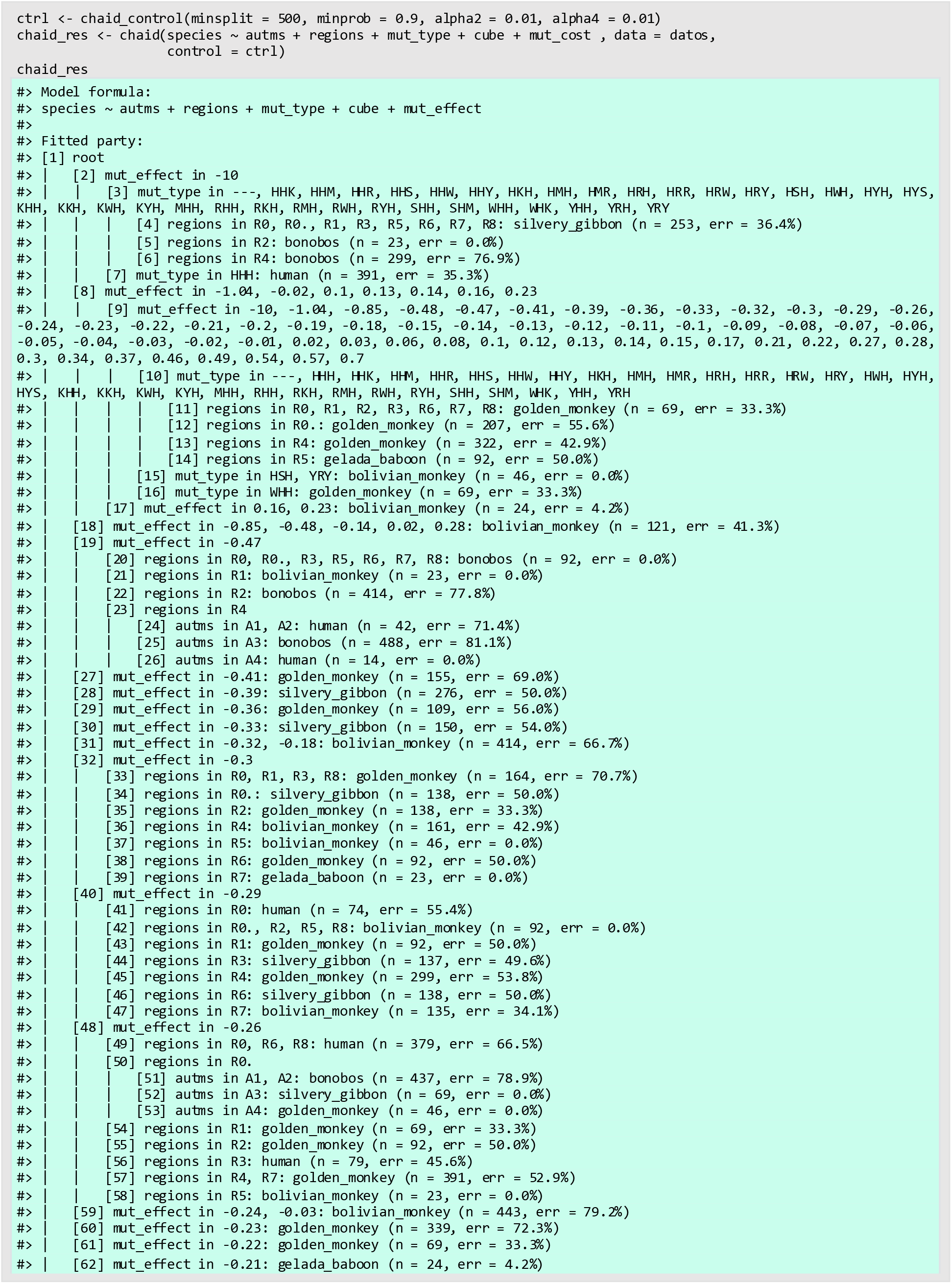
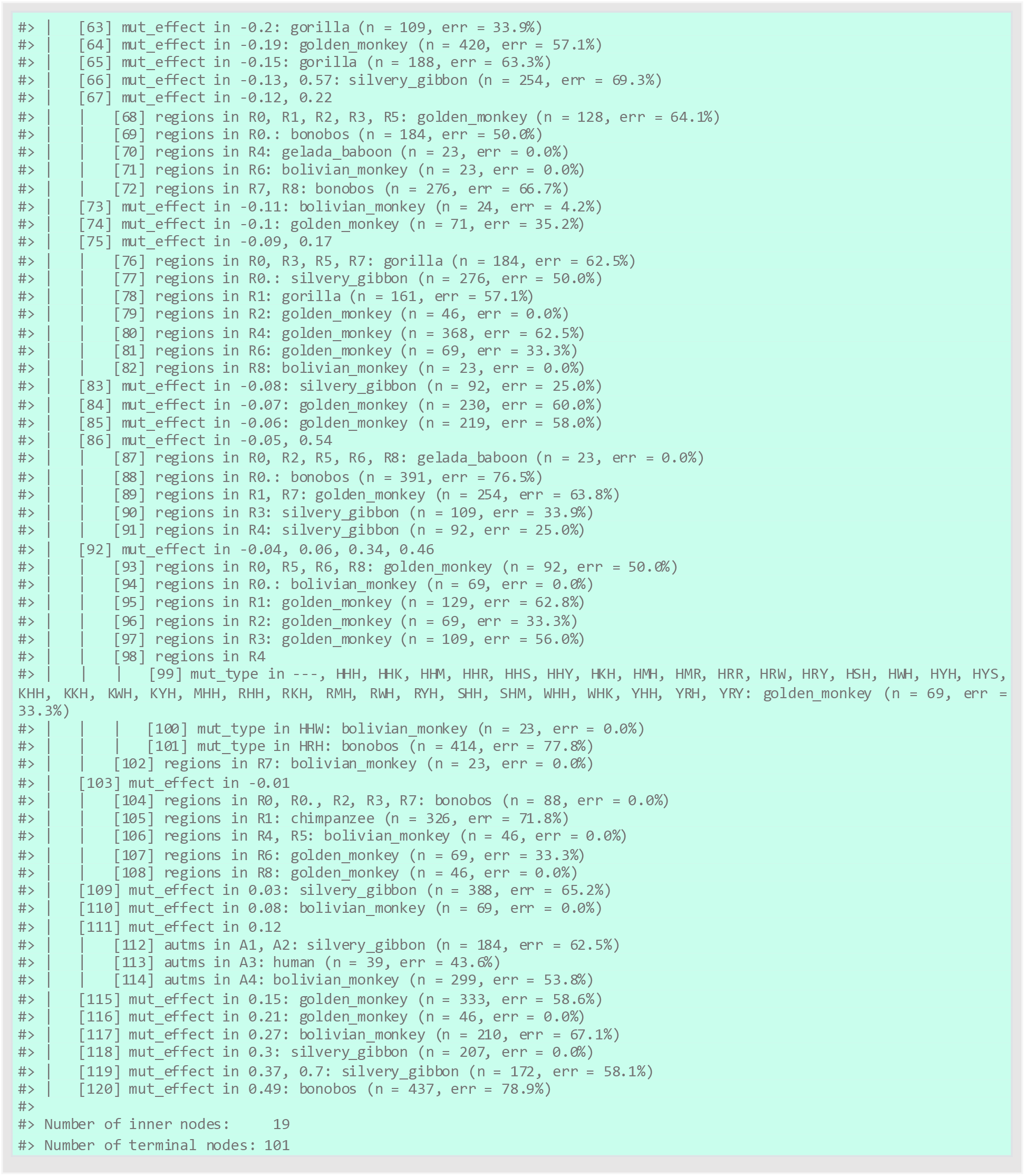

#### 8.2 Using mutational cost in terms of Aminoacid distances

The mutational cost is measured in terms of the average of two aminoacid distance functions, which are based on the codon distance. The entire script is available in the tutorial of *GenomAutomorphism* R package [28]: https://genomaths.github.io/genomautomorphism/articles/automorphism_and_decision_tree_II.html

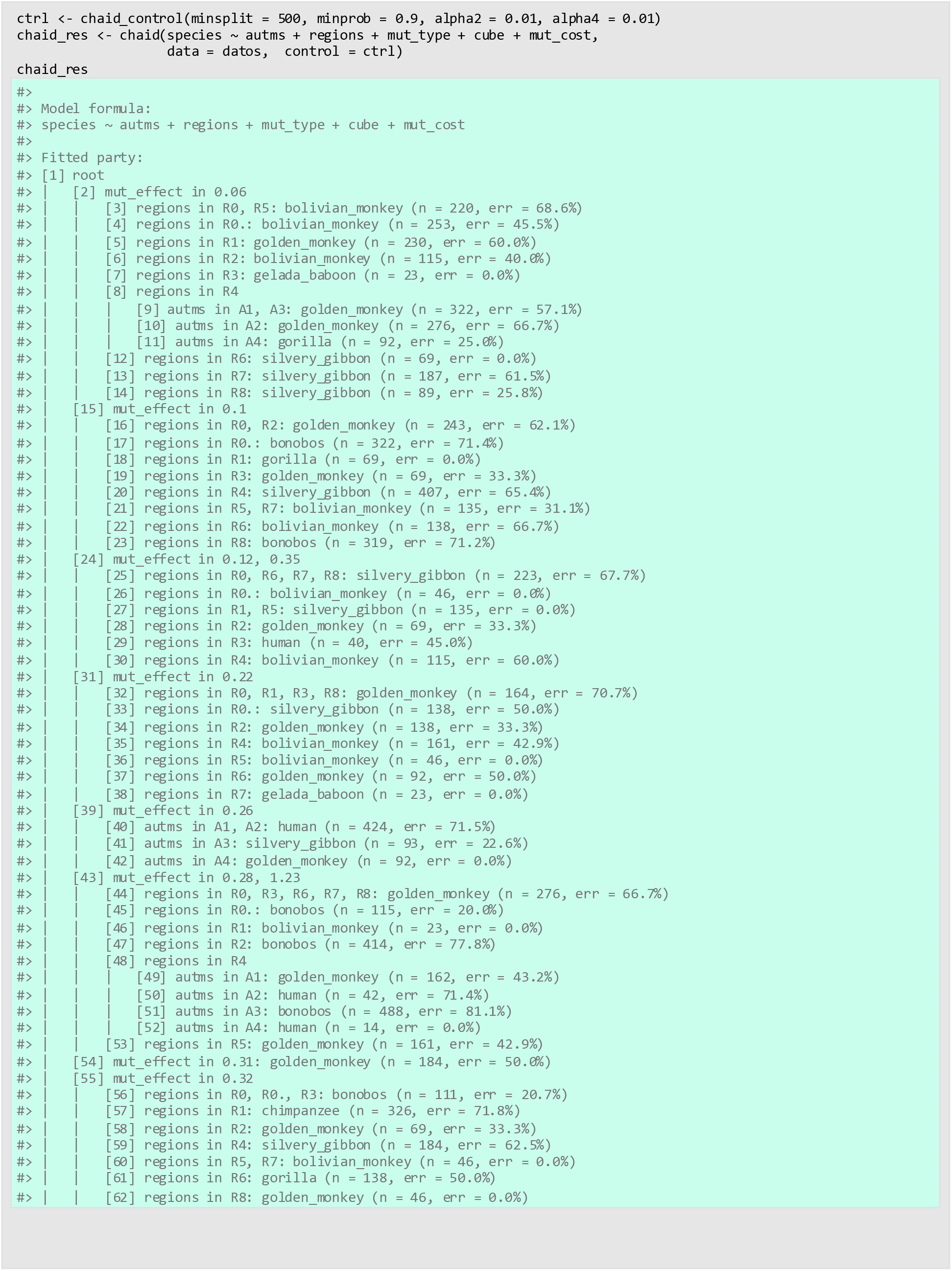
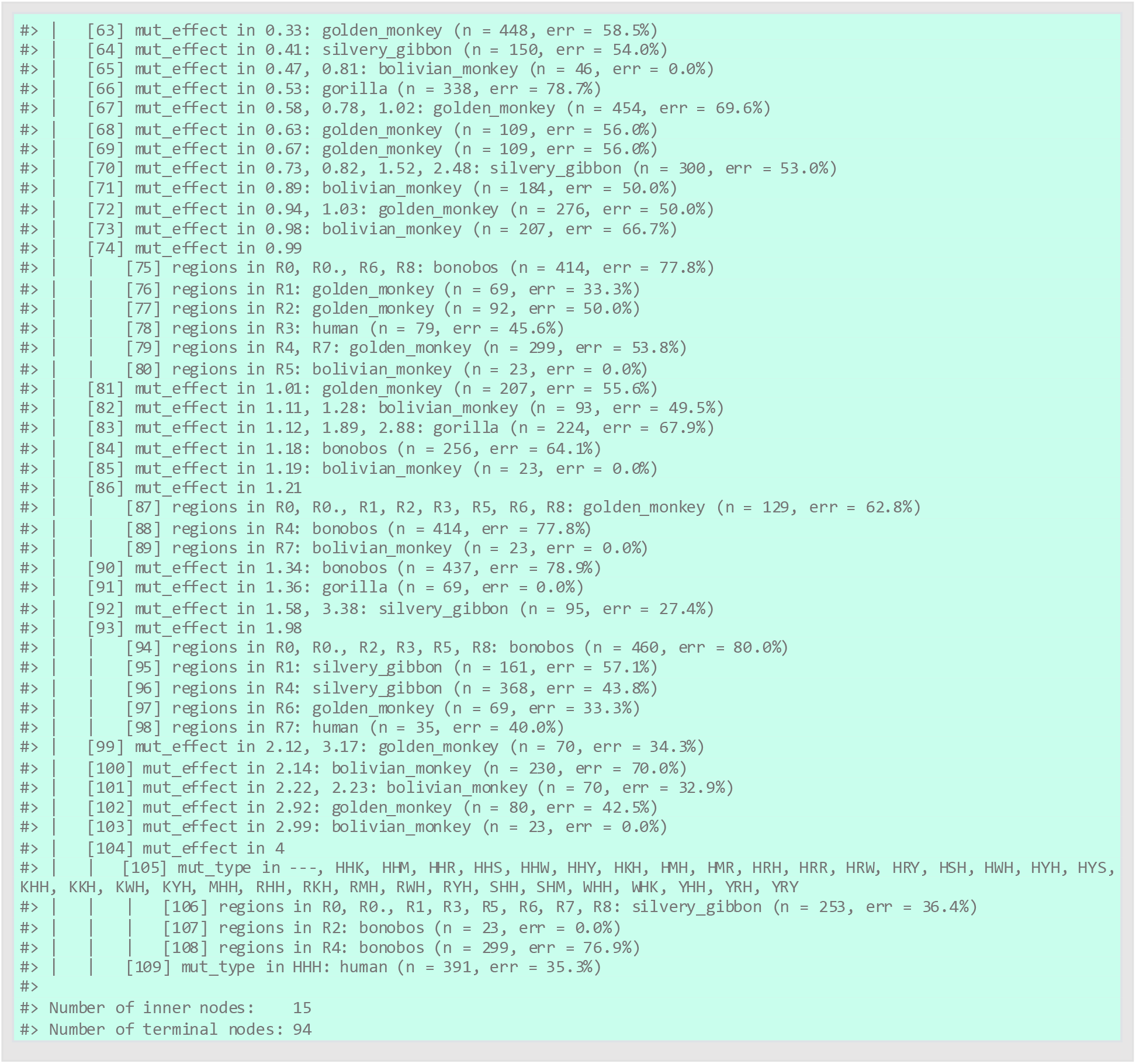

